# HMGN5 escorts oncogenic STAT3 signaling by regulating chromatin landscape in tumorigenesis of breast cancer

**DOI:** 10.1101/2022.01.04.474868

**Authors:** Jiahui Mou, Meijun Huang, Feifei Wang, Xiaoding Xu, Hanqi Xie, Henglei Lu, Mingyang Li, Yu Li, Weiwen Kong, Jing Chen, Ying Xiao, Yiding Chen, Chaochen Wang, Jin Ren

**Affiliations:** Center for Drug Safety Evaluation and Research, State Key Laboratory of Drug Research, Shanghai Institute of Materia Medica, Chinese Academy of Sciences, 501 Haike Road, Shanghai 201203, China.; University of Chinese Academy of Sciences, No.19A Yuquan Road, Beijing 100049, China.; ZJU-UoE Institute, Zhejiang University School of Medicine, International Campus, Zhejiang University, 718 East Haizhou Rd, Haining, Zhejiang 314400, China.; Department of Breast Surgery, the Second Affiliated Hospital, Zhejiang University School of Medicine, 300 Yuan-Ju Rd, Hangzhou, Zhejiang 310009, China.; Nanjing University of Chinese Medicine,Nanjing 210023, China; School of Chinese Materia Medica, Nanjing University of Chinese Medicine, Nanjing 210023, China.; Central Lab of Biomedical Research Center, Sir Run Run Shaw Hospital, School of Medicine, Zhejiang University, Hangzhou, Zhejiang 310020, China.

**Keywords:** HMGN5, STAT3, chromatin landscape, tumorigenesis, breast cancer

## Abstract

Epigenetic alterations are widely linked with carcinogenesis, therefore becoming emerging therapeutic targets in the treatment of cancers, including breast cancer. HMGNs are nucleosome binding proteins, which regulate chromatin structures in a cell type- and disease-specific manner. However, the roles of HMGNs in the tumorigenesis of breast cancer are less known. In this study, we report that HMGNs are highly expressed in 3D-cultured breast cancer cells. HMGN5, a member of HMGNs, controls the proliferation, invasion and metastasis of breast cancer cells in vitro and in vivo. Clinically, HMGN5 is an unfavorable prognostic marker in patients. Mechanistically, HMGN5 is governed by active STAT3 transcriptionally and further escorts STAT3 to shape oncogenic chromatin landscape and transcriptional program. Lastly, we provide evidence that interference of HMGN5 by nanoparticle-packaged siRNA is potentially an effective approach in breast cancer treatment. Taken together, our findings reveal a novel feed-forward circuit between HMGN5 and STAT3 in promoting breast cancer tumorigenesis and suggest HMGN5 as a novel epigenetic therapeutic-target in STAT3- hyperactive breast cancer.

## Introduction

Breast cancer (BC) is the leading malignancy in women. Worldwide, breast cancer accounts for about 30% of female cancers, and has a mortality-to-incidence ratio of 15% (1). Breast cancer is clinically classified into three major subtypes by the expression of hormone receptor (ER and PR) and HER2 (ERBB2): luminal ER-positive and PR-positive, which is further subdivided into luminal A and B; HER2-positive; and triple-negative breast cancer (TNBC). Accordingly, targeted therapies have been developed for different subtypes. Endocrine therapy for hormone receptor-positive cancer works well, and HER2 antibody is widely used for HER2-positive breast cancer. Without explicit molecular signature, it is more difficult to develop targeted therapy for TNBCs, expect for cases bearing definite genetic aberrations. For TNBCs bearing BRCA1/2 mutations, Poly (ADP-ribose) polymerase (PARP) inhibitors are promising agents in clinic application (2). Besides, TNBCs with PTEN mutations were most likely to benefit from AKT inhibitors (3). In addition to genetic aberrations, epigenetic alterations of DNA methylation, histone modifications and chromatin structures also significantly contribute to breast cancer progression (4). Therefore, the causative epigenetic modulators are also efficient targets for therapeutic interventions. For instance, BRCA1 is frequently silent due to DNA methylation and chromatin inaccessibility in TNBCs, while a combination of SGI-110 and MS275, inhibitors of DNA methyltransferase and histone deacetylase, showed a high anti-tumor ability against TNBC (5). Meanwhile, Bromodomain and Extra-Terminal Domain (BET) family of proteins are readers of acetylated histones that are involved in carcinogenesis. Currently, a new BET inhibitor, ZEN-3694, is being evaluated in TNBC patients without germline mutations of BRCA1 or BRCA2 in clinical trials (6). However, comprehensive understanding of the roles of epigenetic modulators and exploration of novel epigenetic-targets for therapy in breast cancer are still largely needed.

Tissue homeostasis is regulated by biochemical signals generated by dynamic reciprocal cell-cell interactions (7). While 2D cultures do not completely recapitulate the three-dimensional (3D) organization of cells within tissues and organs, 3D-culture systems are known to resemble the in vivo behavior of many cell types as they are able to reproduce the specific biochemical and morphological features similar to the corresponding tissue in vivo (8–10). Likewise, tumor development is shaped by cell-cell contact and microenvironment in vivo. While 2D-cultured cells are widely used in cancer studies, they might fail to phenocopy tumor cells in vivo. As a result, potential therapeutic targets may be neglected in these studies (9, 11). Therefore, 3D cultures, such as the well-established spheroid culture system, better reflect the in vivo behavior of cells in tumor tissues (12–14). By performing proteomic analysis, here we report that HMGN5, a member of High-mobility group nucleosome-binding domain (HMGN) protein family, is highly expressed in 3D-cultured TNBC cells.

HMGNs are a group of nucleosome-binding proteins, widely present in mammals and most vertebrates (15). HMGNs destabilize linker histone H1 binding to nucleosomes by competing for chromatin binding sites, unfolding higher-order chromatin structures and modulating transcription (16, 17). In addition, HMGNs mediate several histone modifications, by which they regulate the chromatin architecture at the nucleosome level (16). HMGN5 is one of the five members in the HMGNs family, whose protein can quickly move into the nucleus and interact with nucleosomes, thereby affecting the transcriptional process. Previous studies (18) have indicated that HMGN5 is associated with a range of human cancers. However, the precise function of HMGN5 in breast cancer is obscure. In this study, we demonstrate a critical role of HMGN5 in proliferation, invasion and tumorigenesis of breast cancer both in vitro and vivo. Analysis of clinical samples showed that HMGN5 expression was markedly upregulated during breast cancer progression and correlated with poor prognosis. Additionally, HMGN5 escorted STAT3-mediated oncogenic signaling and is a promising candidate for epigenetic therapy.

## Materials and Methods

### Cell culture and reagents

All cell lines in this study were obtained from the American Type Culture Collection (ATCC, USA). MDA-MB-231, MDA-MB-468 and MDA-MB-453 were grown in L15 medium (Invitrogen, USA) supplemented with 10% fetal bovine serum (Gibco) at 37 °C in air. The BT549, T47D, MCF7, MCF-10A, HEK293 and HEK293T cells were incubated at 37 °C with 5% CO2. BT549 and T47D were grown in RPMI1640 (Gibco, USA) with 10% fetal bovine serum. MCF7, HEK293 and HEK293T were grown in DMEM (Hyclone, USA) supplemented with 10% fetal bovine serum. MCF-10A was cultured with MEBM medium (Lonza, Switzerland) supplemented with 10% fetal bovine serum. All cell lines were authenticated by STR and tested for mycoplasma contamination.

### Tumorsphere formation

Single-cell suspension was prepared and MDA-MB-231 and T47D cells in the medium were seeded in ultra-low attachment dishes to form tumorspheres. Tumorspheres were maintained in DMEM/F12 (Gibco), supplemented with B27 serum replacement (Thermo Fisher Scientific, USA, 17504044), 20 ng/ml basic FGF (sino biological, China, 10014-HNAE), 20 ng/ml recombinant human EGF (sino biological, China, 10605-HNAY), heparin (MedChemExpress, China, HY-17567A) for up to 7 days.

### Mass spectrometry (MS) analysis

MDA-MB-231 tumorsphere and MDA-MB-231 adherent cells were lysed in SDT lysis buffer. Mass spectrometry was performed in Shanghai Institute of Materia Medica, Chinese Academy of Sciences following the standard protocol. Protein quantification was based on TMT-label quantitative analysis.

### RNA extraction and real-time qPCR (RT-qPCR)

Total RNA was extracted by TRIzol reagent (Invitrogen) from cells and reversed transcribed into cDNA using PrimeScript RT Master Mix (TaKaRa, Japan). qPCR was performed to detect mRNA expression of genes by TB Green PCR Kit (TaKaRa) and primers are shown in table S2.

### Western blotting and co-immunoprecipitation (co-IP)

For western blotting, anti-STAT3(9139s), anti-p-STAT3 Y705(9145s), anti-GAPDH (97166) were obtained from Cell Signaling Technology. Anti-HMGN5 (PA5- 50468) was purchased from Invitrogen. Anti-FLAG (F3156) was purchased from Sigma Aldrich.

For co-IP, the clarified supernatants were first incubated with anti-FLAG (Sigma Aldrich, F3156) or anti-HMGN5 antibody overnight at 4 °C, and then incubated with Dynabeads protein G beads (Invitrogen, 10004D) for 2h at 4 °C, and the precipitates were washed five times with RIPA and analyzed by western blotting.

### Cell proliferation and colony formation assay

For the cell proliferation assays, 5,000 cells were seeded into 96-well plates and cell viability was assessed by the Cell Counting Kit-8 (CCK-8) (Dojindo, Japan) at 72h after transfection.

For the colony formation assay, 1,000 cells were seeded into 6-well plates for 10 days. Then, the cells were fixed with 4% paraformaldehyde, washed with PBS, stained with 0.05% crystal violet and washed with water. Finally, the dye was eluted with 1mL of 33% acetic acid, and the optical density (OD) was detected at 595 nm with the microplate reader.

### Animal experiments

All animal experiments were carried out according to the Institutional Ethical Guidelines on animal care and were approved by the Institutional Animal Care and Use Committee, Shanghai Institute of Materia Medica, Chinese Academy of Sciences (2020-03-RJ-212, 2021-05-RJ-240). All mice were housed in a temperature-controlled pathogen-free environment on a 12h light/dark cycle, and had ad libitum access to food and water. In all trials, 4–6-week-old female BALB/c nude mice were purchased from JH-LabAnimal in China.

To explore the effect of HMGN5 on breast tumor growing, each mouse was injected subcutaneously with 2 × 106 MDA-MB-231 cells that stably expressed sh- control or sh-HMGN5. Tumor volume was measured every 3 days after injection. Four weeks later, mice were sacrificed, and then the xenografts were collected for weight mensuration and gene expression analysis.

For in vivo metastasis analysis, 25 mice were randomly allocated to sh-control and sh-HMGN5 group. 2 × 106 MDA-MB-231 cells stably expressing luciferase and sh- control or sh-HMGN5 were injected into the tail vein of mice. Bioluminescence imaging was performed using the IVIS Lumina II (Caliper LS) every week. For luminescence monitoring, the mice were anesthetized using 2% isoflurane, followed by D-luciferin (J&K, China) injection according to the manufacturer’s protocol. Eight weeks after cell injection, the lungs of mice were collected for counting the pulmonary metastatic nodules. Mice bearing over 20 % weight loss and poor condition were euthanized earlier according to “AVMA Guidelines for the Euthanasia of Animals: 2020 Edition”. The lungs of all mice were fixed in formalin and processed with paraffin embedding and Hematoxylin and Eosin (H&E) staining.

For PS@PEI-siRNA treatment experiment, each mouse was injected subcutaneously with 2 × 106 MDA-MB-231 cells. When tumors reached a volume of about 50 mm3, the animals were randomized into treatment and control groups of 7 mice per group. HMGN5 siRNA loaded into polymersomes was injected intravenously at 4.5mg/kg twice a week. The tumors were measured every 3 days for 3 weeks, and the tumor volume (mm3) was calculated using the following formula: V=1/2× length× width2. After 3 weeks, the mice were sacrificed, and the tumor tissues were obtained for weight mensuration and gene expression analysis.

### Migration and invasion assays

The cell invasion and migration assays were performed using the Transwell Permeable Supports with 8μm pore (Corning, USA, 3422). Briefly, 1.5 × 105 cells/cm2 were seeded into apical chambers with the basement membrane pretreated with (for cell invasion assay) or without (for cell migration assay) 20μg Matrigel (Corning, 354248). After 24 h, the transwell was fixed with 4% PFA for 15 min and stained with hematoxylin for 15 min. The cells on the upper side of basement membrane were gently removed by cotton swabs, while the cells on the lower side of basement membrane were imaged. ImageJ was then used to calculate the number of transferred cells.

### Human BC TMA

Tissue microarray (TMA) containing 161 tissue spots from 152 breast cancer patients was obtained from Shanghai Outdo Biotech Co.,Ltd. The TMA was stained with antibodies against p-STAT3 Y705 (Abcam, ab76315) and HMGN5 (Invitrogen, PA5-50468). Positive signal was graded by two automatized histology quantification program (Definiens Tissue Studio® 4.0 or ZEN).

### Immunohistochemistry staining (IHC)

Five-micrometer-thick paraffin sections of human BC patients were obtained from the Second Affiliated Hospital of Zhejiang University. Antigen repair was performed for 10 min in the citric acid buffer (PH 6.0). Then, the treated sections were incubated with primary antibodies against HMGN5 or p-STAT3 at 4 °C overnight. HRP conjugated Goat anti-Rabbit lgG (Active Motif, USA, 15015) and DAB Kit (Zhongshanjinqiao, China, ZLI9018) was then used for immunochemical reaction. IHC score was assessed by pathologists according to the percentage of stained cells (0–4) and staining intensity (0–3).

### RNA-Seq and analysis

Briefly, mRNA was enriched with Oligo-dT magnetic beads followed by reverse transcription, fragmentation and library construction. Sequencing was conducted on an BGISEQ at BGI-Wuhan (China). For RNA-Seq analysis, raw reads were trimmed and filtered first. Then filtered reads were aligned to the GRCh38 human reference transcriptome using salmon (v1.4.0) for count matrix generation (57). Differentially expressed genes were identified using R package DESeq2 (v1.32.0) (58). The genes with adjusted p-value <0.05 and fold change >1.5 were considered as differentially expressed genes. GSEA was calculated by R package fgsea (v1.18.0) (59) and gene sets of hallmark gene sets and curated gene sets were obtained from MSigDB (https://www.gsea-msigdb.org/gsea/msigdb ). Batch correction was performed by ComBat_seq function in R package sva (v3.40.0) when needed(60).

### CUT&Tag and data processing

50,000 cells were harvested and the CUT&Tag assay was performed using a commercial kit (Vazyme, China, TD901-TD902) according to the manufacturer’s protocol. Briefly, cells were collected and washed followed by incubation with Concanavalin A-coated magnetic beads. Next, cells were incubated with anti-Acetyl- Histone3(K27) (Cell Signaling Technology) primary antibody for 2h, and then 1h with secondary antibody at room temperature. Hyperactive pG-Tn5 transposonase was then added to perform tagmentation. The reaction was terminated and DNA fragments were extracted by PCI, followed by PCR amplification using indexed P5 and P7 primers. Purified libraries were then sent for paired-end sequencing on NovaSeq 6000 (Illumina).

The STAT3 ChIP-Seq data were obtained from GEO database (GSM4608989). ChIP-Seq reads were aligned to human reference genome GRCh38 using Bowtie2 (61). Then the STAT3 binding sites were identified by peak-calling using MACS2 (v2.2.7.1) (62) with q-value 0.05. The H3K27ac Cut&Tag sequencing reads were filtered and trimmed before alignment. H3K27ac enriched peaks were then called using MACS2 (v2.2.7.1) with broadpeak option on. H3K27ac peak density were calculated by annotatePeaks.pl function in HOMER2 (63). The STAT3 ChIP-Seq and H3K27ac Cut&Tag tracks were visualized in IGV browser (64).

### Immunofluorescence (IF) staining

Cells treated with IL6 (sinobiological 10395-HNAE) for 15min were fixed in 4% paraformaldehyde for 5 min and then fixed in methanol stored at -20 °C for 10min. Next, cells were permeabilized with 0.2% Triton X-100 for 5 min. After blocking with 1% BSA in PBS for 1 h, the cells were then incubated with primary antibodies overnight at 4°C and then the secondary antibody conjugated with Alexa Fluor 488 (green) (Thermo Fisher Scientific, 35502) or Alexa Flour 594 (red) (Invitrogen, A24207). Nuclei were stained with DAPI (Beyotime, China). A confocal microscope (Leica-SP8) was used for image capture.

### Plasmid construction

The FLAG-STAT3 and FLAG-HMGN5 were purchased from OBiO (China). The FLAG-STAT3C plasmids were constructed as described previously(65). The double strand DNA coding HMGN5 shRNA and STAT3 shRNA were synthesized by Sangon Biotech (Shanghai, China) and cloned into the lentiviral vector pLKO.1-Puro (Addgene, 8453).

### RNAi treatment

30nM synthesized siRNAs were transfected with Lipofectamine RNAiMAX (Invitrogen) according to the manufacturer’s instructions. The oligonucleotide sequences targeting STAT3 and HMGN5 mRNA are shown in table S3.

### The luciferase reporter assay

The promoter region and mutant of HMGN5 was cloned into pGL4.10 luciferase reporter vectors. HEK293 cells were first transfected with siRNA (30nM) for 24h, and then transfected with the reporter constructs (0.5μg) in 24-well plates. SV40promoter- Renilla plasmids were co-transfected as internal control. Forty-eight hours after transfection, cell extracts were prepared and the luciferase activity was measured by the Dual-Luciferase Reporter Assay System (Promega, USA). The primers used for the cloning HMGN5 promoter and its mutant are listed below.

Primers for HMGN5 promoter: 5′-GTAAACGTGCTGCTCTACGACATGA-3′ and 5′-AGCTCCAACTGAAGGTCCCTGA-3′. Primers for HMGN5 promoter mutant: 5′-GGAAAGTCTAAGAGGGCGAGGAGAAGA-3′ and 5′-GGAATCGTCAAAAATCAATGTTTCAACAGATCT-3′

### Chromatin immunoprecipitation (ChIP) assay

MDA-MB-231 cells were cross-linked by 1% formaldehyde for 10 min at room temperature. ChIP assay was performed using the ChIP Assay Kit (Cell Signaling Technology, 9003) according to the manufacturer’s protocol. Anti-STAT3 (9139s) antibody was purchased from Cell Signaling Technology. Primers at predicted STAT3 binding sites were used for qPCR assays.

### Statistics

All in vitro experiments were performed three times independently. Quantitative data from in vitro experiments were shown as mean ± SD; while the data from in vivo experiments were shown as mean ±SEM. One-Way ANOVA test and Student’s two-tailed t-test was used for statistical analysis with multi-group and two-group experiments, respectively. Two-way ANOVA test was used for analyzing the statistical change of tumor volume in tumor xenograft experiments. Kaplan-Meier (KM) survival analysis was performed using survival package in R and the p-value was assessed by log-rank test. p < 0.05 was considered significant statistically.

## Results

### HMGN5 expression is upregulated in 3D-cultured BC cells

Compared to 2D culture, three-dimensional (3D) cell culture mimics the tumor development in vivo, providing a better system to characterize tumor cells and identify therapeutic targets. To reveal the differential molecular profiles of 2D- and 3D-cultured breast cancer cells, we carried mass spectrometry (MS) analysis of both spheres and adherent cells of a typical triple-negative breast cancer cell line MDA-MB-231 (19) (Fig. 1A). As shown in Fig. 1B, 4,587 proteins were detected by MS, among which 1,478 proteins displayed significantly higher levels in 3D condition, while 1,283 proteins were significantly expressed at a lower level. Gene-set enrichment analysis (GSEA) showed that genes associated with cellular metabolisms were significantly enriched in 2D-cultured cells. For example, CRAT (carnitine O-acetyltransferase) and ACSF2 (acyl-CoA synthetase family member 2) are expressed by 3.6- and 2.9-fold higher in 2D-cultured cells respectively, both of which are important enzymes for lipid metabolism (Table S1). These data suggested the cells were more active in metabolism, probably due to easier access to nutrients and oxygen when cultured under 2D condition. Whereas, genes associated with ribosome biogenesis, chromosome organization and chromosome segregation were significantly enriched in 3D-cultured cells, which indicated that the transcription as well as the translation process was dramatically altered in 3D-cultured cells (Fig. 1C and S1A-B). Among the 3D-enriched genes that are associated with chromosome organization, we noticed a nucleosome binding protein family, HMGNs. HMGN family consists of 5 members, HMGN1-5. Interestingly, most of the members were expressed at a higher level in 3D condition according to MS results (Fig. 1B and S1C). Consistently, RT-qPCR results showed that mRNA levels of HMGNs were subtly but significantly increased in 3D condition (Fig. 1D), implying HMGN family members may participate in chromosome re-organization in breast cancer tumorigenesis. Actually, a few studies have reported that HMGN2 and HMGN5 may be involved in breast carcinogenesis (20–22). Recently, a genome-wide CRISPR screen has been performed to identify genes that are involved in cancer cell proliferation (23). We analyzed the screen data in breast cancer cell lines in the study and found that HMGN5 contributes to the proliferation of 8 lines out of the 21 screened breast cancer cell lines. More interestingly, in 6 of 11 basal-type breast cancer cells, HMGN5 was characterized as proliferation-relevant gene. In contrast, HMGN1-4 was barely identified in the screens (Fig. 1E). These data suggested that HMGN5 is possibly a critical gene for breast cancer tumorigenesis, especially in basal-type breast cancers. To further verify that HMGN5 is enriched in 3D-cultured breast cancer cells, we employed western blot analysis of HMGN5 in 2D- and 3D-cultured MDA-MB-231 and T47D (a luminal B type breast cancer cell line) cells. Consistent with our data in MS, HMGN5 was more abundant in 3D-cultured condition in both cells (Fig. 1F). These results suggested that HMGN5 is likely to be involved in breast tumorigenesis in general.

**Fig. 1.**
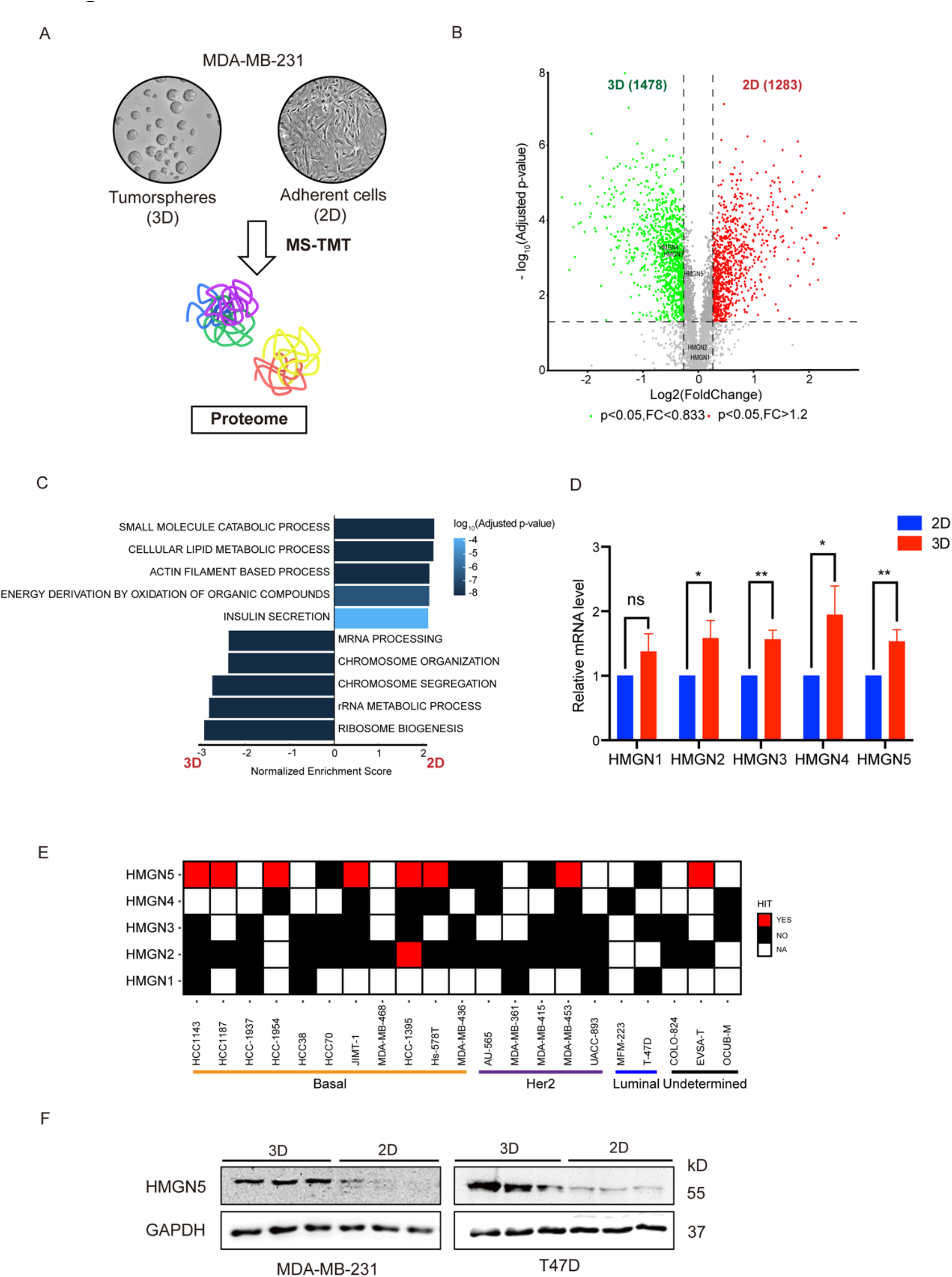
Analysis of differentially expressed genes between 2D- and 3D-cultured MDA-MB-231 cells. (A) Schematic presentation of identifying the differentially expressed proteins between 2D-cultured (adherent cells) and 3D-cultured (tumorspheres) MDA-MB-231 cells. (B-C) Volcano plot (B) of differentially expressed proteins in adherent cells versus tumorspheres of MDA-MB-231 (n=3) detected by Mass spectrometry (MS). HMGNs are also labeled. (C) GSEA of MS data using the hallmark gene sets. (D) HMGNs mRNA levels in adherent cells and tumorspheres of MDA-MB-231 are detected by RT-qPCR. *p <0.05, **p <0.01. (E) Heatmap showing the public CRISPR screen results in breast cancer lines (23). Cell lines are grouped based on molecular subtypes in depmap (https://depmap.org/). HIT score presents if the gene is identified as cell-essential gene. The HIT score is assigned as not applicable (NA), if the gene is not detected in the screen. (F) High expression of HMGN5 in tumorspheres of MDA-MB-231 (left panel) and T47D (Right panel) is confirmed by western blotting when compared with adherent cells.

### HMGN5 promotes BC cell proliferation

To explore the role of HMGN5 in breast cancer, we first examined the function of HMGN5 in the proliferation of breast cancer cells. Three independent siRNAs were used to interfere HMGN5 expression in MDA-MB-231 and T47D cells. The RT-qPCR and western blot results showed that HMGN5 was successfully down-regulated over 85% by either of the siRNAs (Fig. 2A&B). Along with HMGN5 down-regulation, the proliferation rate of MDA-MB-231 cells was significantly decreased (Fig. 2C). Similarly, the growth of T47D, MCF-7 and BT549 cells were all attenuated by HMGN5 knockdown significantly (Fig. 2D, S2A, S2B). These findings were in line with the result of CRISPR screens (23)(Fig. 1E) that HMGN5 is widely required for breast cancer cell proliferation. Notably, the viability of normal breast epithelial cells MCF- 10A was almost unaffected when HMGN5 was knocked down (Fig. 2E), indicating that the role of HMGN5 in proliferation is limited to cancer cells. We further performed colony formation assay in HMGN5 knockdown cells. Intriguingly, with the knockdown of HMGN5, MDA-MB-231 (Fig. 2F and S2C) and T47D cells (Fig. 2G) formed drastically fewer and smaller colonies compared with those in control groups. These data demonstrated that HMGN5 specifically regulates breast cancer cell growth and clonal expansion in vitro.

**Fig. 2.**
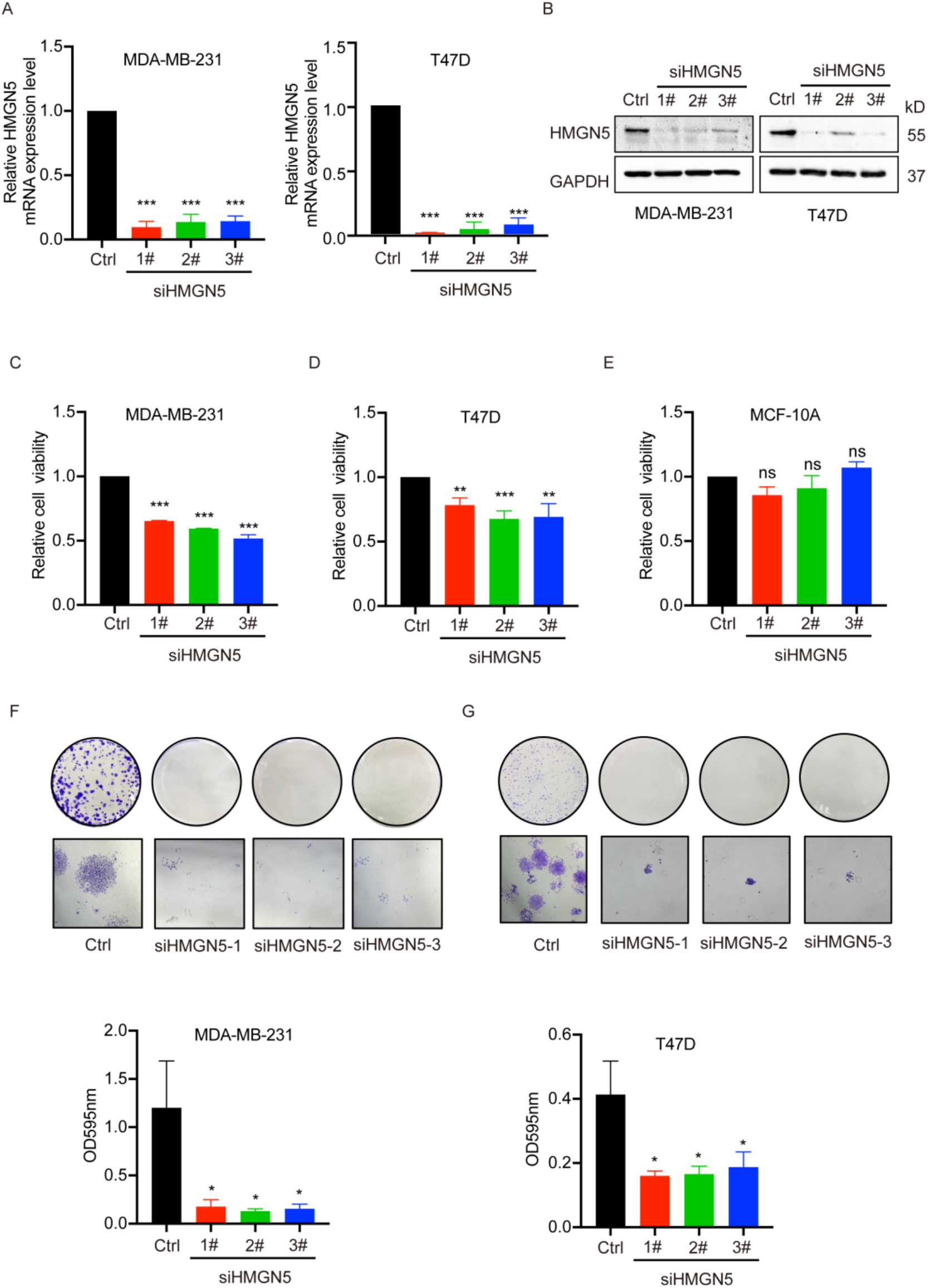
HMGN5 depletion inhibits breast cancer cell growth and clonal expansion in vitro. (A-B) The knockdown efficiency of HMGN5 in MDA-MB-231 and T47D cells is verified by RT-qPCR (A) and western blotting (B). (C-E) Relative cell viabilities of MDA-MB-231 (C), T47D (D) and MCF-10A cells (E) are determined by CCK-8. (F-G) The abilities of clonal expansion of MDA-MB-231 (F) and T47D (G) cells are assessed by colony formation assays. *p <0.05, **p <0.01, ***p <0.001 in one-way ANOVA test.

### HMGN5 controls breast tumor growth

Although HMGN5 regulates breast cancer cell proliferation, the impact of HMGN5 depletion on proliferation was moderate. Given that HMGN5 is highly expressed in 3D-cultured cells, we speculated that HMGN5 plays a more important role in breast cancer tumorigenesis. To verify our speculation, we first evaluated whether HMGN5 was important in the spheroidization of BC cells by tumorsphere formation assay. We stably knocked down HMGN5 by around 95% in MDA-MB-231 cells and 70% in T47D cells with short hairpin RNA (shRNA) (Fig. 3A), and then observed that the number and sizes of tumorspheres were both dramatically decreased (Fig. 3B). These results indicated that HMGN5 is critical for the survival of breast cancer cells under 3D-cultured condition. To further explore the role of HMGN5 in tumorigenesis in vivo, we injected MDA-MB-231 cells infected with short hairpin RNAs targeting HMGN5 (sh-HMGN5) and control cells into immunodeficient mice to form tumor xenografts. We found that the xenografts from sh-HMGN5 cells grew extremely slowly compared to the control group (Fig. 3C). After 30 days, the median weight of tumor xenografts derived from control cells reached around 200mg, while that from sh- HMGN5 was only about 10mg; the size of the tumors was also drastically smaller in sh-HMGN5 group (Fig. 3D&E). We also detected HMGN5 mRNA and protein abundance in the tumor tissues, and HMGN5 was successfully depleted as expected (Fig. 3F&G). In addition, similar results were observed in sh-HMGN5 T47D cells (Fig. 3H-L). These results indicated that the inhibition of HMGN5 severely slowed down the tumor growth of breast cancer in vivo, providing strong evidence that HMGN5 is a key gene in breast cancer tumorigenesis.

**Fig. 3.**
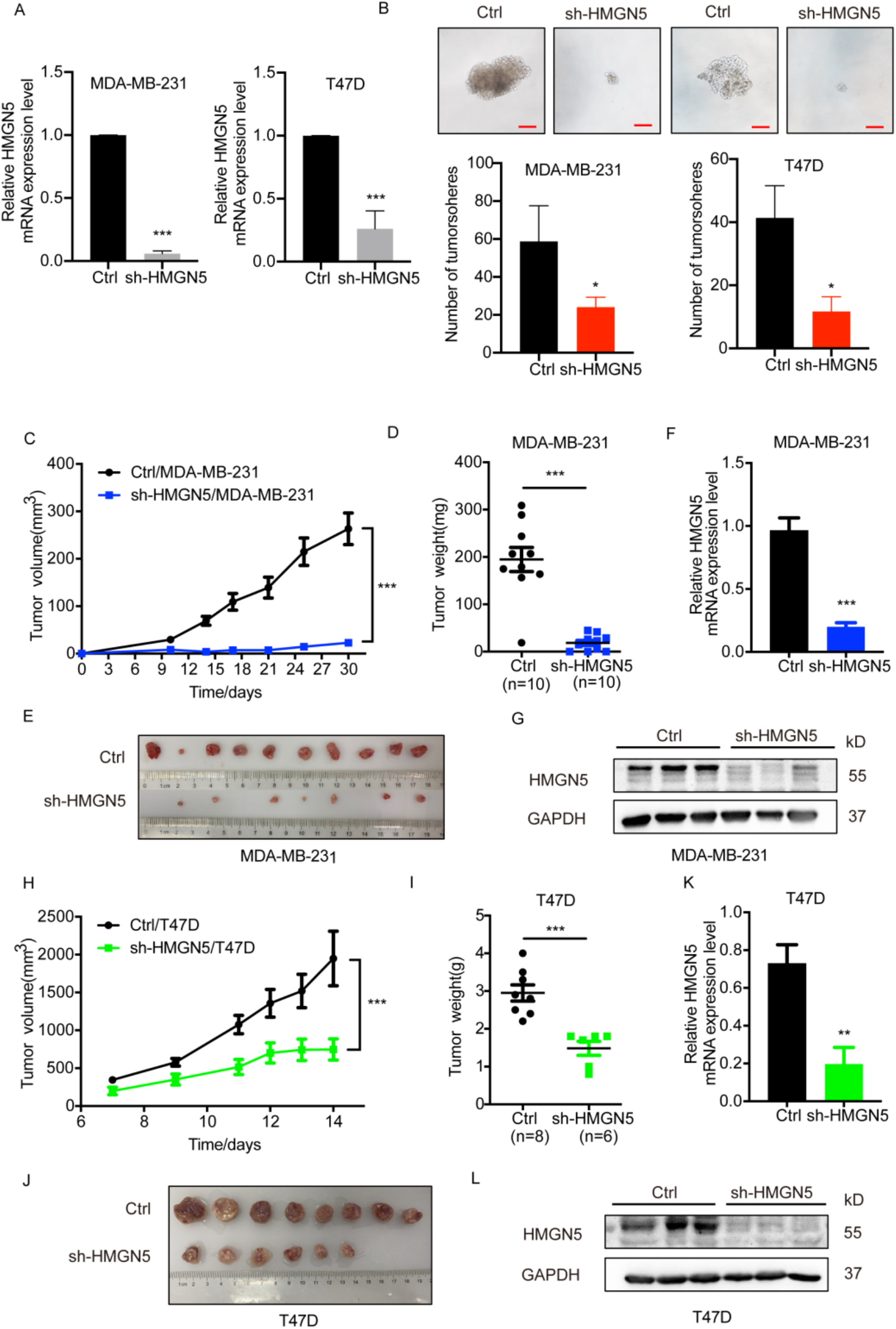
HMGN5 depletion impedes breast tumor growth in vivo. (A) The knockdown efficiency of HMGN5 in MDA-MB-231 and T47D cell lines infected with HMGN5- shRNA lentiviruses are verified by RT-qPCR. (B) Representative images and quantification of tumorspheres formed by control and sh-HMGN5 MDA-MB-231 cells and T47D cell lines. Scale bar, 20 µm. (C) The growth curve of tumors formed from MDA-MB-231 cell line containing sh-HMGN5 or sh-control (n = 10 per group). The size of tumor xenografts in BALB/c nude mice was monitored every 3 days. (D) Barplot presenting the weights of tumors formed from stable MDA-MB-231 cell line containing sh-HMGN5 or sh-control (n = 10 per group). (E) Photographs of tumors in two groups are shown. (F-G) HMGN5 levels in the tumors of two groups are examined by RT-qPCR (F) and western blotting (G). (H-L) Similar experiments are performed in T47D cell line containing sh-HMGN5 or sh-control (n = 10 per group). The tumor volume (H), tumor weight (I), tumor photograph (J), HMGN5 mRNA level (K), and HMGN5 protein level (L) are assessed. For A, B, D, F, I and K, p-value is determined by Student’s two-tailed t-test. For C, H, p-value is determined by two-way ANOVA test. *p <0.05, **p <0.01, ***p <0.001 as indicated.

### HMGN5 facilitates breast tumor metastasis

To further investigate the effects of HMGN5 on mobility and invasiveness in breast cancer, we performed transwell migration assays. It showed that suppression of HMGN5 expression significantly inhibited the migration of MDA-MB-231 cells (Fig. 4A). In addition, silencing HMGN5 led to significant decrease of the invasive ability of MDA-MB-231 cells (Fig. 4B), revealing that HMGN5 also contributed to breast cancer cell migration and invasion. Western blot analysis of Epithelial-mesenchymal transition (EMT) associated proteins showed that ZEB-1 and N-cadherin were strikingly downregulated in sh-HMGN5 cells (Fig. 4C), suggesting that HMGN5 possibly contributes to migration and invasion of BC cells in an EMT dependent manner. Furthermore, to validate the role of HMGN5 in tumor metastasis in vivo, we delivered sh-HMGN5 cells carrying luciferase into immunodeficient mice by tail vein injection. Consistently, 7 of 15 mice in the control group died within 8 weeks, while all the 10 mice injected with sh-HMGN5 cells survived (Fig. 4D). Histological analysis demonstrated that HMGN5 knockdown significantly reduced the incidence of lung metastasis and the number of metastatic lung nodules (Fig. 4E). In addition, whole-body bioluminescence imaging (BLI) was performed to disclose metastatic burdens in the mice. Starting from 5 weeks after injection, apparent metastatic lesions were detected in the lungs of mice injected with control cells. In contrast, metastatic cancer cells were barely observed when HMGN5 was knocked down (Fig.4F and S3A). Moreover, combining BLI and histological analysis demonstrated that over 80% of control mice developed lung metastasis after 8 weeks; on the contrary, only 10% of sh- HMGN5 mice had (Fig.4G). Together, these data manifested the critical function of HMGN5 in breast cancer metastasis.

**Fig. 4.**
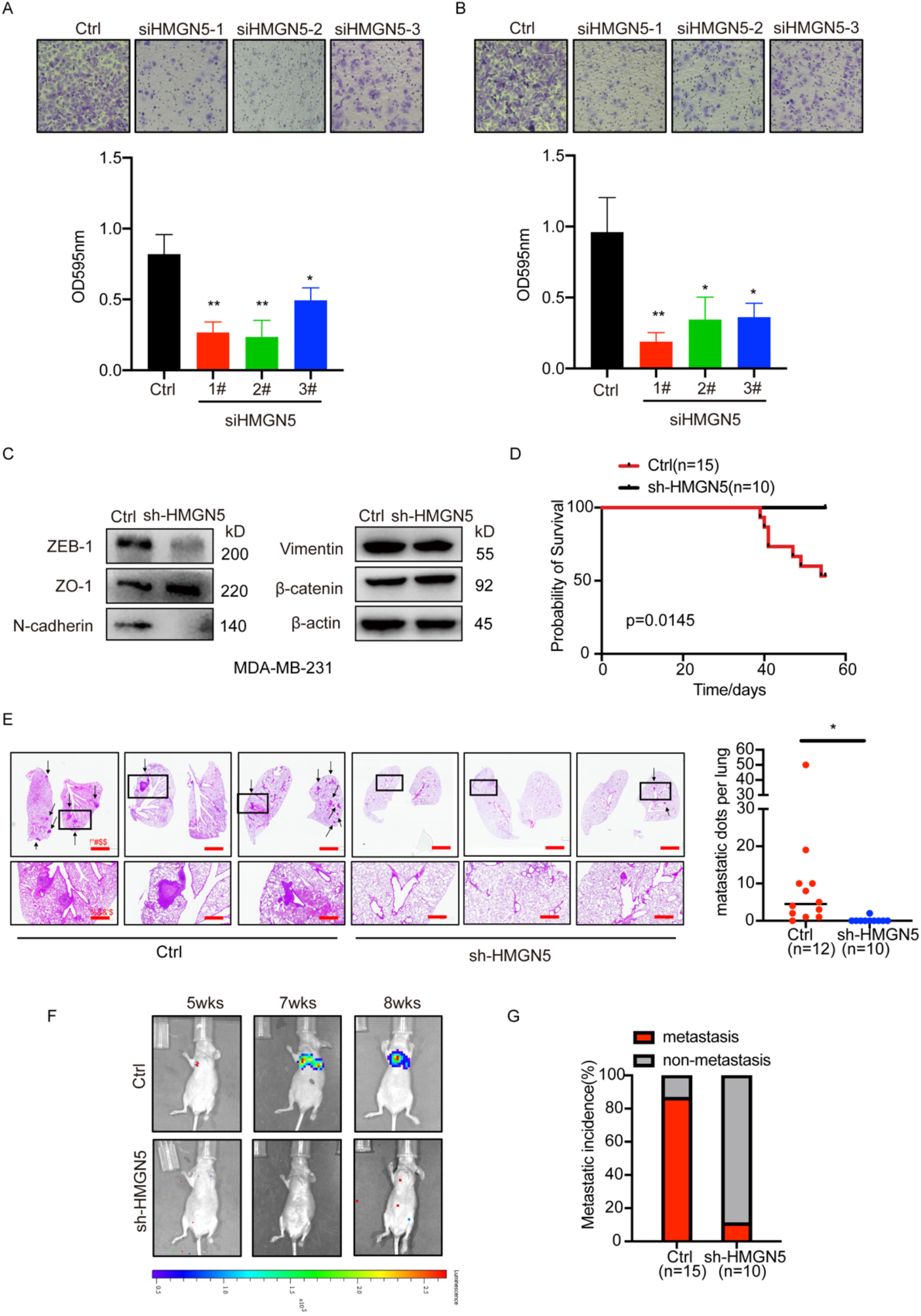
Knockdown of HMGN5 suppresses breast cancer cell metastasis in vitro and in vivo. (A-B) Transwell migration (A) and transwell invasion assays (B) of MDA-MB-231 cells transfected with siHMGN5. Representive images and the number of migrating cells are shown. (C) Expression of EMT markers (ZEB1, ZO1, N-cadherin, vimentin and n-catenin) in HMGN5-deficiency MDA-MB-231 cells are detected by western blotting. (D) The Kaplan-Meier survival curves of mice injected with MDA-MB-231 cells containing sh-HMGN5 (sh-HMGN5 group, n=10) or sh-control (control group, n=15). (E) Representative images of H&E staining of lung sections from mice in the indicated group. The black arrowheads indicated the metastatic nodules. Scale bar, 2500µm (top); scale bar, 800µm (bottom). The number of pulmonary nodules are also calculated as shown on the right (control group, n=12; sh-HMGN5 group, n=10). (F) Representative bioluminescence image of mice in the indicated group at 5, 7, and 8-week after injections of cancer cells. (G) Barplot showing the metastatic incidence of mice in the indicated group (control group, n=15; sh-HMGN5 group, n=10). For A and B, p-value is determined by one-way ANOVA test. For D, p-value is determined by log- rank (Mantel-Cox) test. For E, p-value is determined by Student’s two-tailed t-test. *p <0.05, **p <0.01.

### HMGN5 is an unfavorable prognosis marker in breast cancer patients

To determine the clinical implication of HMGN5, we analyzed the transcriptomic data from 1,081 breast cancer patients in TCGA. High HMGN5 expression was significantly associated with poor prognosis in the patients (Fig. 5A). Moreover, we performed immunohistochemical staining (IHC) analysis of a tissue microarray, which includes a cohort of 88 luminal A, 20 luminal B, 15 HER2-positive, 29 triple-negative breast cancer samples and 9 normal adjacent tissues. According to the H-score of staining density, we grouped the cancer samples into HMGN5-, HMGN5low and HMGN5high classes (Fig. 5B). In general, HMGN5 was highly expressed in all the four subtypes of breast cancer samples compared to normal breast samples (Fig. 5C). And more than 48% of luminal A and HER2-positive samples, and 75% of luminal B and triple-negative subtype were HMGN5high (Fig. 5D). Additionally, the Kaplan-Meier survival analysis showed that HMGN5 expression levels were reversely correlated with the overall survival time of breast cancer patients, which was consistent with the result from the TCGA breast cancer cohort (Fig. 5E). Therefore, these observations indicated that HMGN5 is an unfavorable prognostic marker in clinics.

**Fig. 5.**
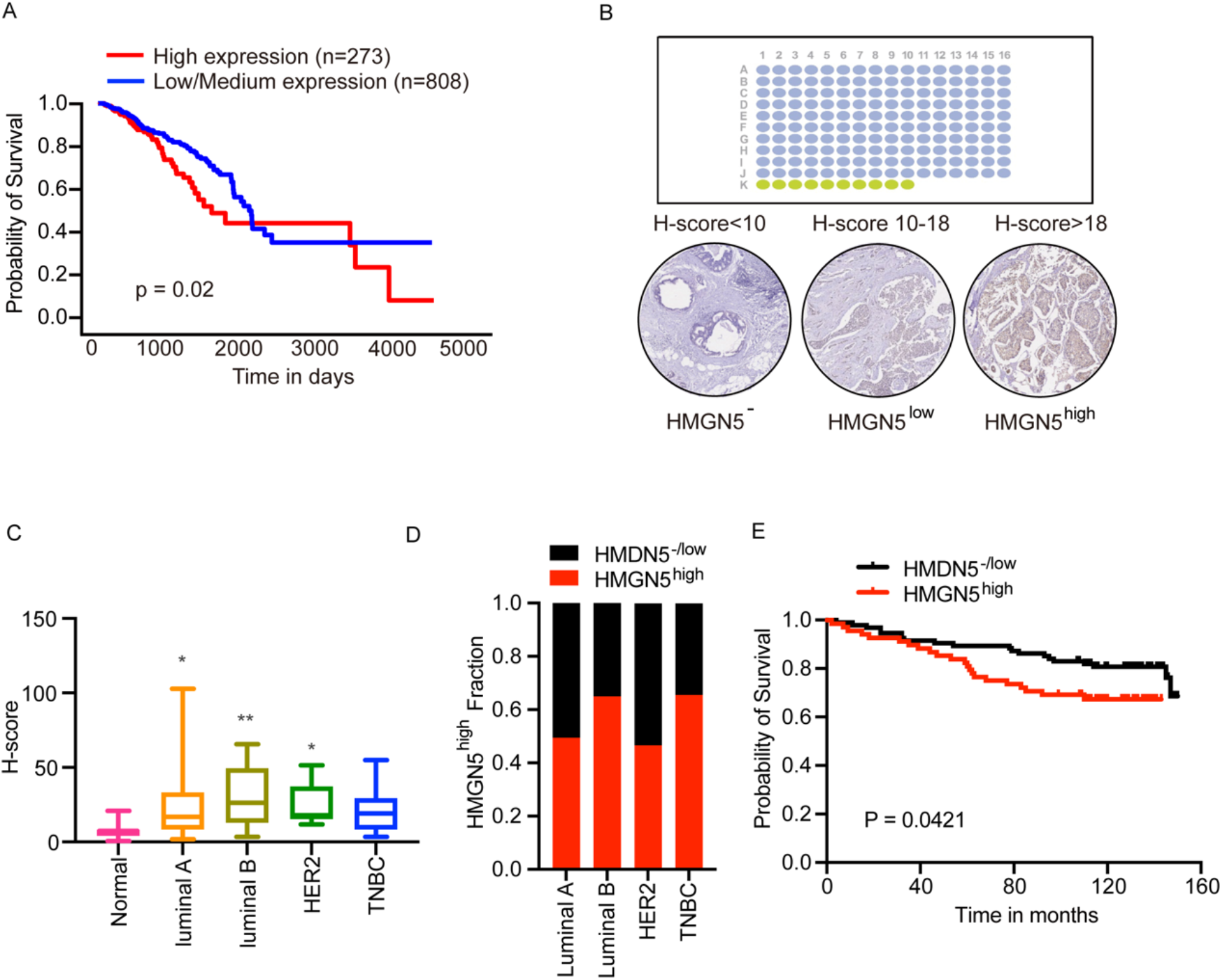
Clinical relevance of HMGN5 in human breast cancer. (A) Kaplan-Meier survival curves stratified by HMGN5 expression in breast cancer patients from the Cancer Genome Atlas (TCGA-BRCA). (B) IHC of HMGN5 in tissue microarray containing152 breast cancers and 9 non-cancerous. Representative images of negative (H-score<10), weak (10<H-score<18) and strong (H-score>18) staining are shown. The median H-score of these 161 cases is 18. (C) HMGN5 protein levels in all the four subtypes of breast cancer samples and normal breast samples in the cohort. (D) The proportions of HMGN5high and HMGN5low cases in the indicated subtype group in our cohort. (E) Kaplan-Meier survival curves stratified by HMGN5 expression in our cohort. For A and E, p-value is determined by log-rank (Mantel-Cox) test. For C, p-value is determined by one-way ANOVA test. *p <0.05, **p <0.01.

### HMGN5 is indispensable for STAT3-mediated oncogenic transcriptome and epigenome in breast cancer

To reveal the underlying molecular mechanism of HMGN5 promoting breast cancer tumorigenesis, we firstly performed RNA-Seq in HMGN5 knockdown MDA-MB-231 cells. Gene enrichment analysis of disease-associated genes in DisGeNET showed that the genes downregulated in siHMGN5 cells were mostly enriched in invasive carcinoma of breast (Fig. 6A), consistent with our findings that siHMGN5 led to attenuated proliferation, invasion and migration of the cancer cells. For instance, CDKN1A and LGR5, known to be involved in breast carcinogenesis, were down- regulated significantly upon HMGN5 interference (Fig. S4A). In addition, gene enrichment analysis of curated gene sets in MSigDB demonstrated that genes associated with EMT were also significantly decreased in siHMGN5 cells, including ZEB1 and N-Cadherin (CDH2) (Fig. S4A&B). These data reproduced our western blot results that the expression of ZEB-1 and N-Cadhenrin were dependent on HMGN5 (Fig. 4C). Intriguingly, GSEA of hallmark genes in MSigDB revealed that IL6/STAT3 signaling was significantly downregulated when HMGN5 was depleted (Fig. 6B and S4C). IL6/STAT3 signaling is a well-studied oncogenic pathway in breast cancer, involved in tumor growth, metastasis and drug resistance (24, 25). It implied that HMGN5 might control breast carcinogenesis through IL6/STAT3 pathway, at least partially. Therefore, we also knocked down STAT3 by siRNAs in MDA-MB-231 cells and compared the transcriptomic changes in siSTAT3 and siHMGN5 cells in parallel. Surprisingly, the majority of genes whose expression was dependent on STAT3 also relied on HMGN5 existence. On the other hand, the genes that were upregulated by STAT3 knockdown were also increased when HMGN5 were knockdown (Fig. 6C). The data suggested that the STAT3-mediated transcriptional program are highly dependent on HMGN5.

**Fig. 6.**
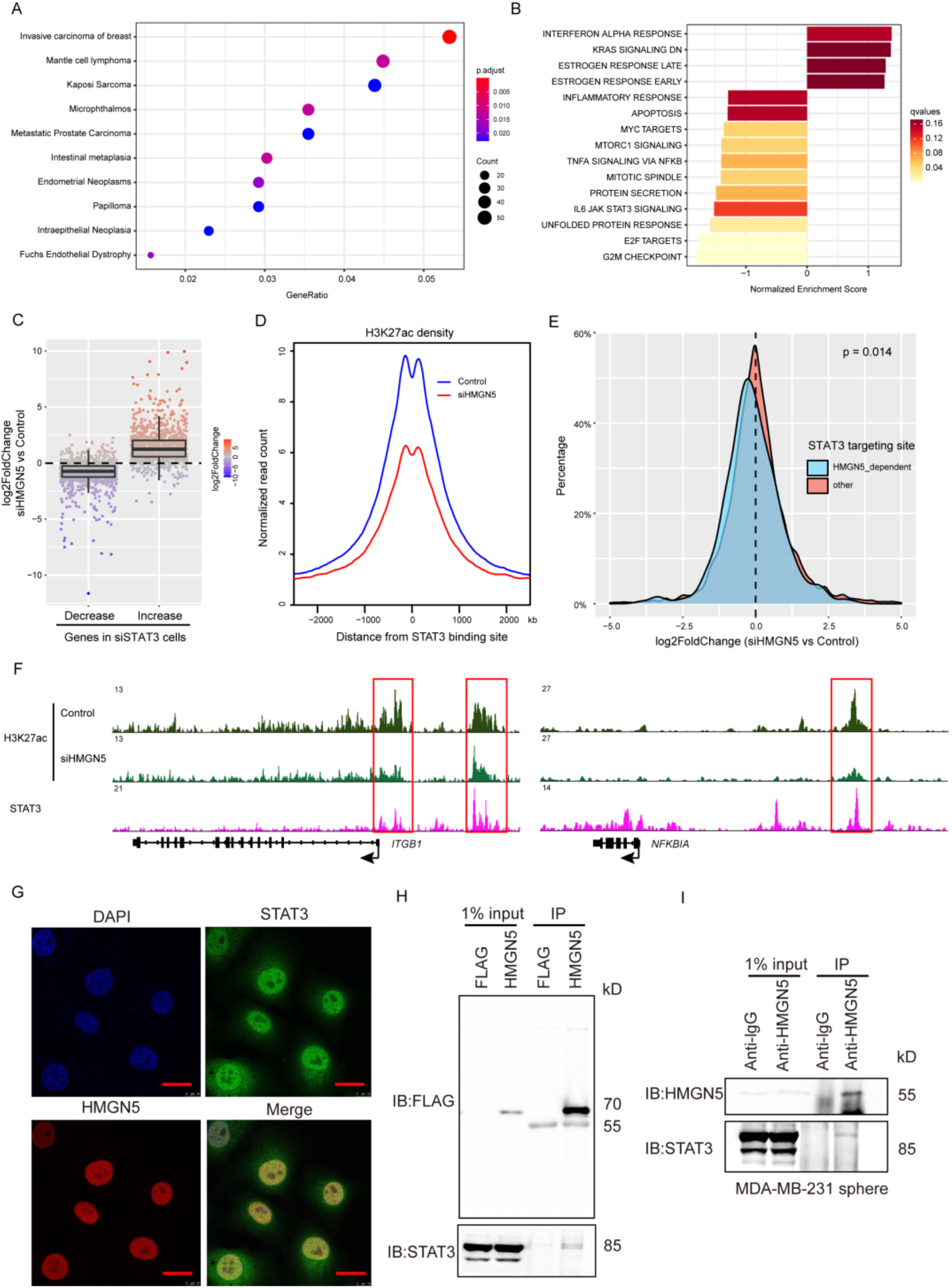
HMGN5 is an epigenetic regulator of STAT3 signaling in breast cancer cells. (A) DisGeNET associated gene enrichment analysis of HMGN5-dependent genes (FC>2, p<0.05) in MDA-MB-231 cells. (B) GSEA of hallmark gene sets in MsigDB based on the rank of expression foldchange in HMGN5 deficient cells vs. control cells. (C) Boxplot of expression changes of genes in siHMGN5 cells. Genes were grouped according to their behavior in siSTAT3 cells. Increase group: expression is significantly increased (FC>2, p<0.05) in siSTAT3 cells compared with control cells; Decrease group: expression is significantly decreased in siSTAT3 cells. (D) Average profiles of H3K27ac density centered at STAT3 binding sites in HMGN5-deficient (siHMGN5) or control MDA-MB-231 cells. (E) Density plot of expression foldchange of STAT3 binding site-associated genes in siHMGN5 cells vs. control cells. Genes were grouped according to the H3K27ac density change of their associated STAT3 binding regions in siHMGN5 cells. The sites with over 2-fold decrease of H3K27ac density in siHMGN5 cells are defined as HMGN5-dependent sites. (F) H3K27ac and STAT3 profiles at the indicated gene loci. (G) Immunofluorescent staining of STAT3 and HMGN5 with 50ng/mL IL6 stimulation for 15min in MDA-MB-231 cells. Scale bars, 20 µm. (H) Co-immunoprecipitation assays (Co-IP) of HEK293 cells were transfected with the FLAG-tagged constructs of HMGN5 or FLAG empty vector. (I) The interaction between endogenous STAT3 and HMGN5 in wild-type MDA-MB-231 tumorspheres is demonstrated by co-IP assays.

Then we wondered how HMGN5 regulates STAT3 signaling in breast cancer. Upon IL6/STAT3 signaling activation, STAT3 undergoes phosphorylation, transports from cytoplasm to nucleus, and then activates or represses downstream targets. It was reported that HMGN5 remodels chromatin by binding to nucleosome and regulates chromatin accessibility (26), which led us to hypothesize that HMGN5 may facilitate STAT3 target gene expression by promoting chromatin accessibility. To test this idea, we firstly identified STAT3 binding sites on the genome of MDA-MB-231 cells based on published ChIP-Seq data (27). Next, we employed CUT&Tag assays to measure the H3K27ac (acetylated H3K27) profiles, an epigenetic mark of active chromatin regions across the genome. Compared to control cells, the average intensity of H3K27ac signal on STAT3 binding sites was markedly decreased in HMGN5 knockdown cells (Fig. 6D), indicating that STAT3-mediated chromatin opening relied on HMGN5 in breast cancer. As expected, the expression of genes nearby HMGN5-dependent STAT3 binding sites was significantly reduced in HMGN5 knockout cells in general (Fig. 6E). These data supported our hypothesis that HMGN5 regulates STAT3 target gene expression by controlling chromatin activation. For example, STAT3 bound to the promoter and several potential enhancers of the gene ITGB1, which was reported to play a role in breast cancer development (28). While in HMGN5 knockdown cells, H3K27ac enrichment was decreased significantly on the promoters and the enhancers (Fig. 6F). In parallel, ITGB1 expression was also attenuated in either siSTAT3 or siHMGN5 cells significantly (Fig. S4D). Similar to IGTB1, NFKBIA was also bound by STAT3 on a distal enhancer, whose activation was also dependent on STAT3 and HMGN5 (Fig. 6F&S4D).

Our data suggested that HMGN5 works synergistically with STAT3 on activating target regions on the chromosome. Immunofluorescence staining of STAT3 and HMGN5 in IL6-treated MDA-MB-231 cells showed that HMGN5 strongly colocalized with STAT3 in the nuclei (Fig. 6G). Plus, the result of co-immunoprecipitation assay in HEK293 cells indicated that ectopic FLAG-HMGN5 physically interacted with STAT3 (Fig. 6H). Consistently, endogenous HMGN5 and STAT3 interacted with each other in MDA-MB-231 forming tumorspheres (Fig. 6I). Together, these results provided strong evidence that HMGN5 and STAT3 conspire to shape an oncogenic chromatin landscape in breast tumors.

### HMGN5 is a direct target of STAT3

We noted that STAT3 was also bound to the promoter region of HMGN5 when we analyzed the ChIP-Seq data in MDA-MB-231 cells (Fig. S5A). Besides, according to our RNA-Seq data, HMGN5 expression was strikingly decreased in STAT3 knockdown cells (Fig. S5B). So we speculated that STAT3 also controls the transcription of HMGN5 via binding to its promoter. Therefore, we cloned the 1kb DNA sequence of the HMGN5 promoter in front of a reporter gene. The results of luciferase reporter assay showed that knockdown of STAT3 by siRNA decreased the reporter activity; whereas ectopic expression of a constitutively active STAT3 mutant, STAT3C (21), increased the reporter activity (Fig. 7A). To further pin down the regulatory element in HMGN5 promoter region, we focused on eight predicted STAT3 binding motifs with the consensus sequence TT(N)4 – 6AA throughout the region. ChIP-qPCR assay in MDA-MB-231 cells indicated that p-STAT3 directly bound to the narrow region between -78∼ +32 bp, which contains one predicted STAT3 binding site, named HMGN5-SIE (Fig. 7B). By luciferase reporter assay, we further investigated the response of the HMGN5-SIE to STAT3. As expected, STAT3 knockdown significantly attenuated transcriptional activity of HMGN5-SIE, but not a mutant isoform. Consistently, wild-type, but not mutant HMGN5-SIE responded to exogenous expression of STAT3C (Fig. 7C).

**Fig. 7.**
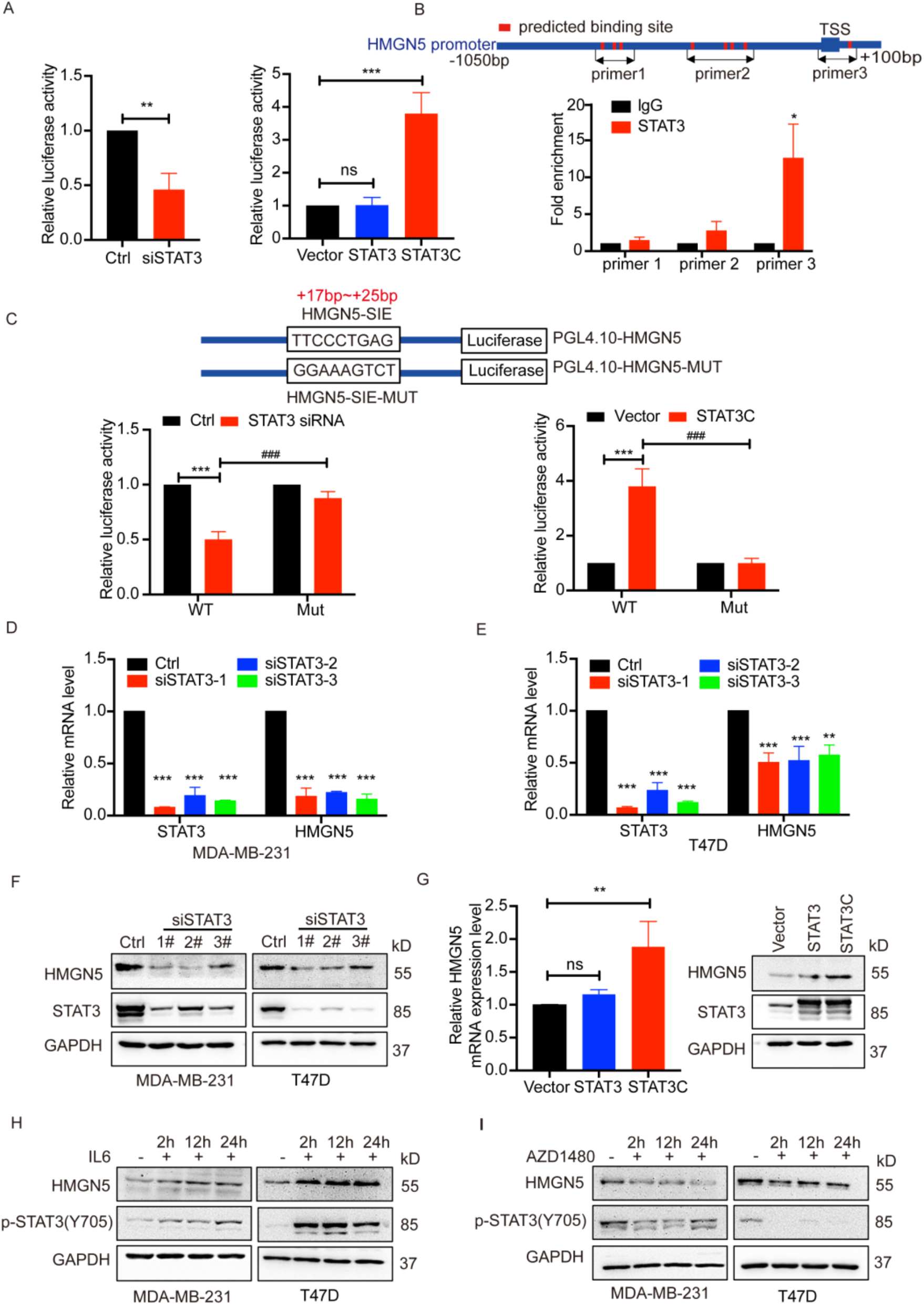
STAT3 transcriptionally activates HMGN5. (A) Effect of STAT3 on the activity of HMGN5 promoter in HEK293 cells is evaluated by reporter assay. STAT3C: constitutive activation of STAT3 (54) (B) ChIP-qPCR assays of STAT3 in MDA-MB- 231 cells. The target regions of indicated primers and the predicted STAT3 binding motifs on HMGN5 promoter are shown in the upper diagram. (C) Schematic diagram of HMGN5 promoter-reporter construct and its mutant. The effects of STAT3 on these two constructs are evaluated by reporter assays. (D-E) STAT3 and HMGN5 mRNA levels in MDA-MB-231 (D) and T47D cells (E) transfected with STAT3 siRNAs are determined using RT-qPCR. (F) HMGN5 and STAT3 protein levels in MDA-MB-231 and T47D cells transfected with STAT3 siRNAs are detected by western blotting. (G) Effects of overexpressing STAT3C on HMGN5 mRNA (left panel) and protein (right panel) levels in HEK293 cells. (H-I) Protein levels of HMGN5 and p-STAT3 were detected at indicated time points by western blotting in MDA-MB-231 and T47D cells treated with 50nM IL6 (H) and 100nM JAK1/2 inhibitor AZD1480 (I). For A left panel, B, p-value is determined by Student’s two-tailed t-test. For A right panel, C, D, E, G, p-value is determined by one-way ANOVA test. *p <0.05, **p <0.01, ***p <0.001, ### p <0.001 as indicated.

To further support the fact of STAT3 controlling HMGN5 expression in breast cancer, we knocked down STAT3 in MDA-MB-231 and T47D cells, in which marked reduction of HMGN5 mRNA levels were observed (Fig. 7D&E). Consequently, the protein level of HMGN5 declined when STAT3 was interfered (Fig. 7F). In contrast, introducing a constitutively active STAT3 mutant, STAT3C (21), into HEK293 cells upregulated HMGN5 mRNA and protein level (Fig. 7G). Moreover, IL6-mediated STAT3 phosphorylation augmented HMGN5 expression in MDA-MB-231 and T47D cells (Fig. 7H). Concurrently, abrogation of STAT3 activation by a Jak2 inhibitor, AZD1480, led to apparent reduction of HMGN5 protein level (Fig. 7I). In addition to MDA-MB-231 and T47D cells, HMGN5 expression was also impaired by STAT3 interference in MCF-7 and the other three breast cancer cells (Fig. S6A-D), revealing that the regulation of HMGN5 by STAT3 universally existed.

### The oncogenic function of HMGN5 is largely through STAT3 signaling

Our results suggested that HMGN5 works synergistically with STAT3 and promotes breast carcinogenesis. However, it is unclear whether the oncogenic function of HMGN5 is solely through the oncogenic STAT3 pathway in tumorigenesis. To answer the question, we over-expressed HMGN5 in the STAT3 knockdown MDA-MB- 231 cells. As expected, downregulation of STAT3 hindered tumor growth of inoculated MDA-MB-231 cells in nude mice; whereas this reduction was slightly reversed by exogenous HMGN5 expression (Fig. 8A-C). Meanwhile, RNA-Seq analysis demonstrated that HMGN5 overexpression barely altered the transcriptomic profile of STAT3 knockout cells (Fig.8D). Among the genes whose expression was significantly associated with STAT3, only a few were rescued by ectopic HMGN5, as shown in the heatmap in Fig. 8E. Altogether, these data suggested that HMGN5 act largely through the STAT3 signaling pathway, although HMGN5 may also participate in other oncogenic signaling.

**Fig. 8.**
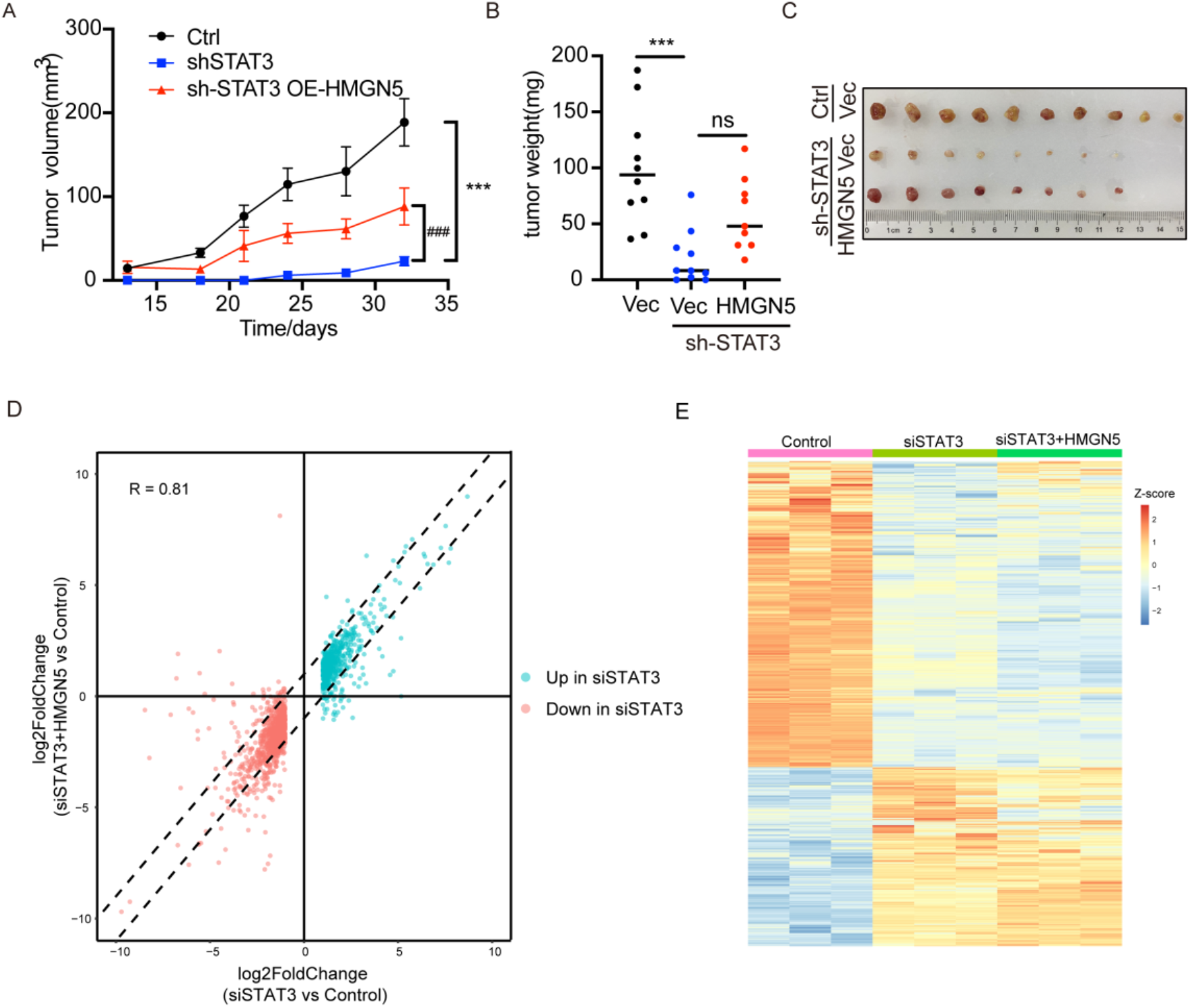
HMGN5-mediated tumorigenesis requires STAT3. (A-C) Effects of HMGN5 overexpression on growth (A) and weights (B) of tumors formed from MDA-MB-231 cells expressing sh-STAT3 (n=10 per group), and the photograph of tumors (C) is also shown. (D) Scatter plot of expression foldchanges of differentially expressed genes in siSTAT3 cells. The R-value of foldchange correlation is also shown. (E) Heatmap of expression levels of differentially expressed genes in siSTAT3 cells (FC > 2, p < 0.05) in the indicated samples. For A, p-value is determined by two-way ANOVA test. For B, p-value is determined by one-way ANOVA test. *p <0.05, **p <0.01, ***p <0.001, ### p<0.001 as indicated.

### HMGN5 is a potential therapeutic target in STAT3-hyperactive breast cancer

Our results indicated that HMGN5 is both a co-activator and a direct target of STAT3 in breast cancer cells. We inquired whether the correlation between the two proteins occurs commonly in breast cancer development. Comparing the HMGN5 and p-STAT3 protein levels of MDA-MB-231 cells in 2D- and 3D-culture conditions, we found that the p-STAT3 levels are remarkably higher in 3D-cultured cells, in accompany with higher HMGN5 levels. More importantly, tumor xenografts derived from the cells also exhibited more abundant p-STAT3(Y705) and HMGN5 than those in 2D condition. Similar phenomena were also observed in T47D cells (Fig. 9A). Together with our finding that HMNG5 is a direct target of STAT3, it suggested that STAT3 signaling is enhanced, thus activating HMGN5 during breast cancer tumorigenesis. In consistent with the result, IHC analyses of p-STAT3 and HMGN5 revealed a high correlation between HMGN5 and p-STAT3 expression in 30 cohorts of human breast cancer specimens in our hands (Fig. 9B). Moreover, cellular colocalization of HMGN5 and p-STAT3 was detected, given to the cellular heterogeneity in the tumor (Fig. 9C).

**Fig. 9.**
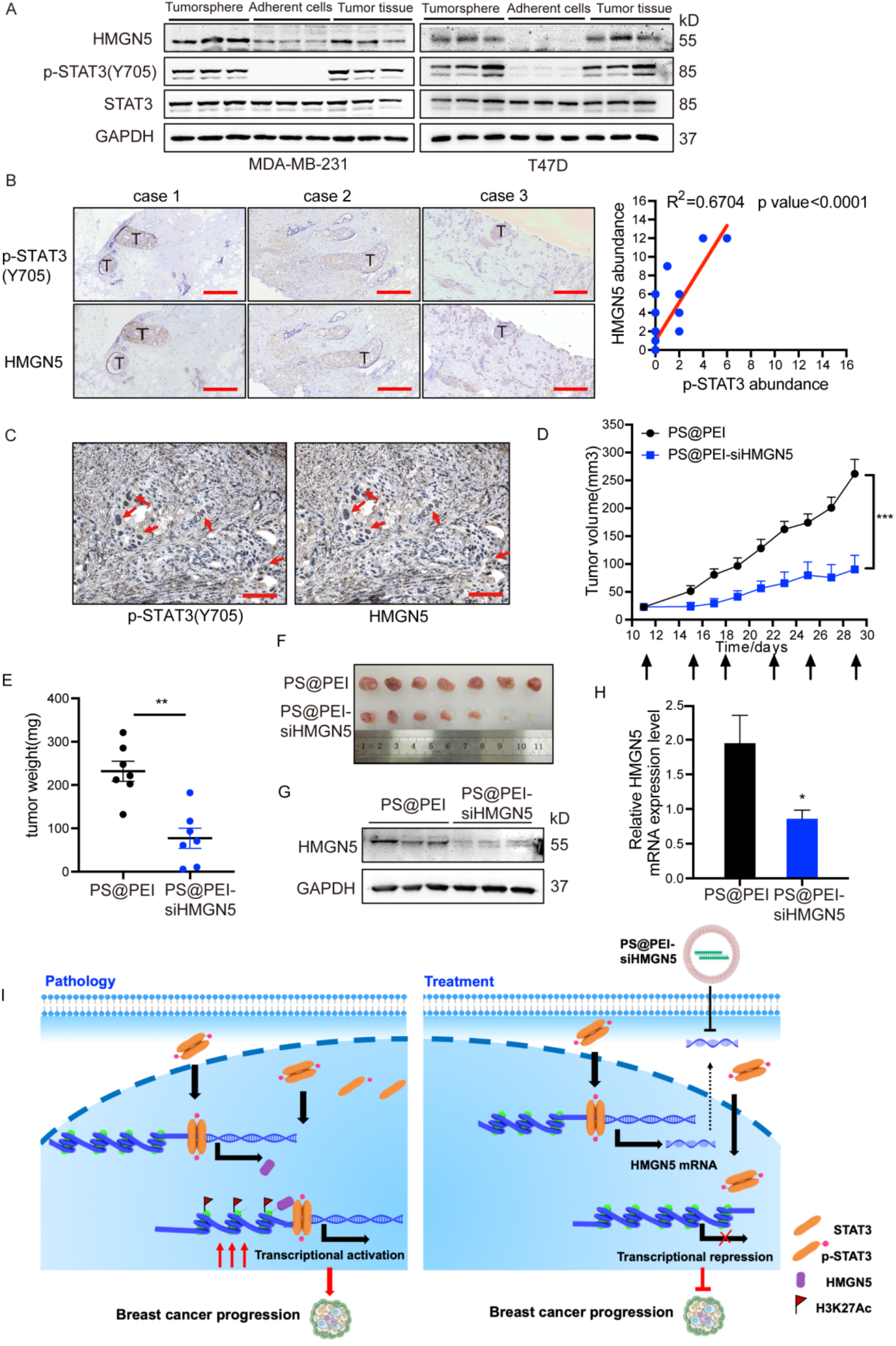
HMGN5 is a therapeutic target in STAT3-hyperactive breast cancer. (A) HMGN5 and p-STAT3 levels are detected by western blotting in adherent cells, tumorspheres and tumor-xenografts of MDA-MB-231 (left panel) and T47D (Right panel). (B) Left panel: representative images of IHC staining of HMGN5 and p-STAT3 in the breast cancer tissues from patients. Scale bar, 500 µm. Right panel: correlation analysis between HMGN5 and p-STAT3 density in patient samples (n=30). (C) Representative images of the cellular colocalization of HMGN5 and p-STAT3 by IHC staining. Scale bar, 100 µm. (D) Effects of PS-PEI-siHMGN5 on tumor growth (n=7 per group). The size of tumor xenografts is monitored every 3 days. (E) Effects of PS- PEI-siHMGN5 on the tumor weights (n=7 per group). (F) Photographs of the tumors in the indicated groups are shown. (G-H) HMGN5 levels in the tumors of indicated groups are examined by western blotting (G) and RT-qPCR (H). (I) A schematic model demonstrating the mechanism of HMGN5 interacting with STAT3 oncogenic signaling in breast cancer and the potential treatment approach targeting HMGN5.The hyper- activated STAT3 signaling triggers the entry of excessive p-STAT3 dimers into the nucleus, which boosts the transcription of HMGN5. HMGN5 further partners with STAT3 and promotes the chromatin remodeling of STAT3-targeted sites, leading to the establishment of oncogenic transcriptome in breast tumorigenesis. For B, p-value is measured by Pearson’s correlation test. For D, p-value is determined by two-way ANOVA test. For E and H, p-value is determined by two-tailed Student’s t test. *p <0.05, **p <0.01, ***p <0.001.

Given to the vital role of HMGN5 in oncogenic STAT3 signaling, it suggested that HMGN5 is a novel epigenetic target to treat STAT3-hyperactive breast cancer, thus urging us to explore a potential approach in clinical application. Specifically, we used the modified virus-mimicking chimeric polymersomes to deliver HMGN5 siRNA to tumor cells to achieve HMGN5 gene silencing (29). The carrier prevented the naked siRNA from being degraded by the ribozyme (RNase), allowing the siRNA to enter the target cell intact, which could greatly improve the therapeutic effect. The anti-tumor efficacy of the HMGN5 siRNA loaded in polymersomes (PS@PEI-siHMGN5) was investigated in vivo. Mice bearing MDA-MB-231 xenografts were treated with the PS@PEI-siHMGN5 or PS@PEI-control for 3 weeks. Marked inhibition of tumor growth was observed in PS@PEI-siHMGN5 treated group (Fig. 9D-F). Analysis of the tumor grafts showed that HMGN5 mRNA and protein levels were reduced by half with the administration of PS@PEI carried siRNA (Fig. 9G and H). Collectively, our data not only reinforced the notion that HMGN5 is a valuable therapeutic target in breast cancer, but also demonstrated that delivery of siRNAs via PS@PEI could be a practical approach to treat HMGN5-high breast cancer clinically.

## Discussion

In conclusion, our findings identified HMGN5 as a novel prognostic biomarker and a potential therapeutic target in breast cancer. As reported by King and Francomano in 2001 (30), the HMGN5 gene is upregulated and confers oncogenic effects in various tumors, such as bladder cancer (31), renal cancer (32), colorectal cancer (33), esophageal squamous cell carcinoma (34) and pancreatic ductal adenocarcinoma (35), etc. HMGN5 appears wildly involved in cancer cell growth, apoptosis, angiogenesis, migration, invasion, and EMT to promote tumorigenesis (36–40). However, few reports revealed the precise molecular mechanisms of HMGN5 in tumorigenesis. In this study, we report a positive regulatory circuit in breast cancer tumorigenesis, by which activated STAT3 enhanced HMGN5 that in turn facilitated STAT3 transcriptional activity in breast cancer (Fig. 9I).

Notably, excessive STAT3 activation and resulting carcinogenesis were also present in cancers besides breast cancer (41, 42). Therefore, it is possible that the interplay between STAT3 and HMGN5 may orchestrate tumorigenesis universally. More interestingly, we found that the activity of the STAT3-HMGN5 circuit was strikingly enhanced in 3D-cultured cells and tumor xenografts compared with 2D- cultured cells, making which an attractive therapeutic target in solid tumor treatment.

Although we showed that the STAT3-HMGN5 circuit was hyperactive in tumor, why it was activated in vivo was not clear. It was known that hypoxic and aberrant microenvironment contribute to the reprogramming of cancer cells in vivo. The level of IL6 was low under normal culture conditions (21% oxygen), but hypoxia (1% oxygen) treatment induced a significant increase in IL6 secretion (43). Thus, STAT3- HMGN5 signaling may be activated under hypoxic conditions along with the activation of IL6. Moreover, YAP signaling pathway for sensing mechanical pressure is reported to regulate STAT3 activity (44). Plus, the resulting alterations in glycolysis and glutaminolysis pathways are frequently found in tumors (45). STAT3 signaling can be constantly activated in the glycolysis process in cancer metabolism (46). Also, extracellular glutamine is reported to activate STAT3, which is necessary and sufficient to mediate the proliferative effects of glutamine on glycolytic and oxidative cancer cells (47). Additionally, STAT3 is one of the representative redox-sensitive transcription factors whose activity is affected by the concentration of reactive oxygen species (ROS). The oxidation and glutathionylation of specific cysteine residues under conditions of excessive oxidative stress impair the DNA-binding and transcriptional activity of STAT3; whereas mild ROS production, along with tyrosine phosphorylation of STAT3, promotes nuclear translocation of STAT3(46). Therefore, the activation of STAT3- HMGN5 circuit could be driven by the above factors together in tumor development.

In this study, we show that HMGN5, among HMGNs, is the major coordinator of STAT3 in breast cancer tumorigenesis. It is worth noting that the other members of HMGNs also participate in breast cancer development. Previous studies indicated that HMGN2 affects STAT5 chromatin accessibility and regulates STAT5 binding and gene transcription in breast cancers (20). HMGN3 has been found upregulated in breast cancer cell lines (48). Although it is unknown if HMGN4 is related to breast cancer, it is reported that HMGN4 promotes thyroid tumorigenesis (49) and hepatocellular carcinoma (50). Further studies are needed to explore the differential roles of HMGNs in breast cancer and the underlying mechanisms.

Given to the important roles of STAT3 in breast cancer (51), different approaches of repressing hyperactive STAT3 have been explored to fight breast cancers. For example, JAK kinase inhibitor, EGFR inhibitor, STAT3 direct inhibitor, STAT3-binding inhibitors, Oligonucleotides (STAT3 decoy), SHP-1 activators, etc. have been developed to treat STAT3-hyperactive cancers (52–54). However, the efficacy of the compounds is not ideal. One of the reasons was that the STAT3 could escape from the inhibition by bypassing JAK signaling or undergoing constitutive active mutation. Targeting HMGN5, the epigenetic modulator of STAT3-dependent oncogenic program, would avoid the above problems, at least partially. Indeed, interference of HMGN5 led to dramatic inhibition of STAT3-mediated oncogenic program in our hands. Therefore, targeting HMGN5 instead of STAT3 per se could be an alternative approach to treat STAT3-hyperactive breast cancers.

Therapeutic interventions targeting epigenetic regulators have attracted attention for years. Chemical inhibitors targeting DNA methyltransferases (DNMT1, DNMT3), histone deacetylases (HDACs), histone methyltransferases (i.e., EZH2, DOT1L) are approved by the US Food and Drug Administration (FDA) in cancer treatment; while drugs targeting histone demethylases (i.e., LSD1), bromodomain proteins (i.e., BETs) are also developed and under clinical investigation (55). Here, we propose that HMGN5 is also a promising epigenetic target in breast cancer therapy. However, small molecules specifically inhibiting HMGNs are not available yet. We proved that delivery of interfering RNA of HMGN5 by nanoparticles might be a feasible approach in clinical application, although the efficacy is not optimal (Fig. 9I). Interestingly, mutation analysis revealed that a novel carbazole, SH-I-14, disrupted the STAT3–DNMT1 interaction, led to the re-expression of tumor-suppressive genes through DNA demethylation, and showed a high anti-proliferative effect in TNBC models (56). Thus, the combination of DNMT inhibitor and HMGN5 interference may give out a better outcome in the treatment of STAT3-hyperactive breast cancer. Nevertheless, an optimized therapeutic strategy requires further investigation.

## Author contributions

Study design, data acquisition and analysis: J.M., C.W., M.H., H.X., M.L., J.R.

Animal experiments, data acquisition and analysis: F.W., X.X., Y.L.W.K

Data analysis of animal histology: H.L.

Clinical samples and data collection: Y.C.

Writing—original draft: J.M.

Writing—review & editing: C.W., J.C., Y.X.

## Acknowledgments

We thank Prof. Zhiyuan Zhong (Soochow University) for preparing the si- HMGN5 loaded virus-mimicking chimaeric polymersomes. We thank the the Institutional Technology Service Center of Shanghai Institute of Materia Medica, Chinese Academy of Sciences for technical assistance in mass spectrometry experiments and analysis. We thank Prof. James Q Wang (Zhejiang University) for reading our manuscript. Especially, we would like to thank the contributors for providing breast cancer tissue specimens. The protocol was approved by the Human Ethics Committee of The Second Affiated Hospital Zhejiang University School of Medicine (2021-0526), and written informed consent was obtained from all participants.

