## Supplementary material for "HMGN5 escorts oncogenic STAT3 signaling by regulating chromatin landscape in tumorigenesis of breast cancer": Fig.S1-S7

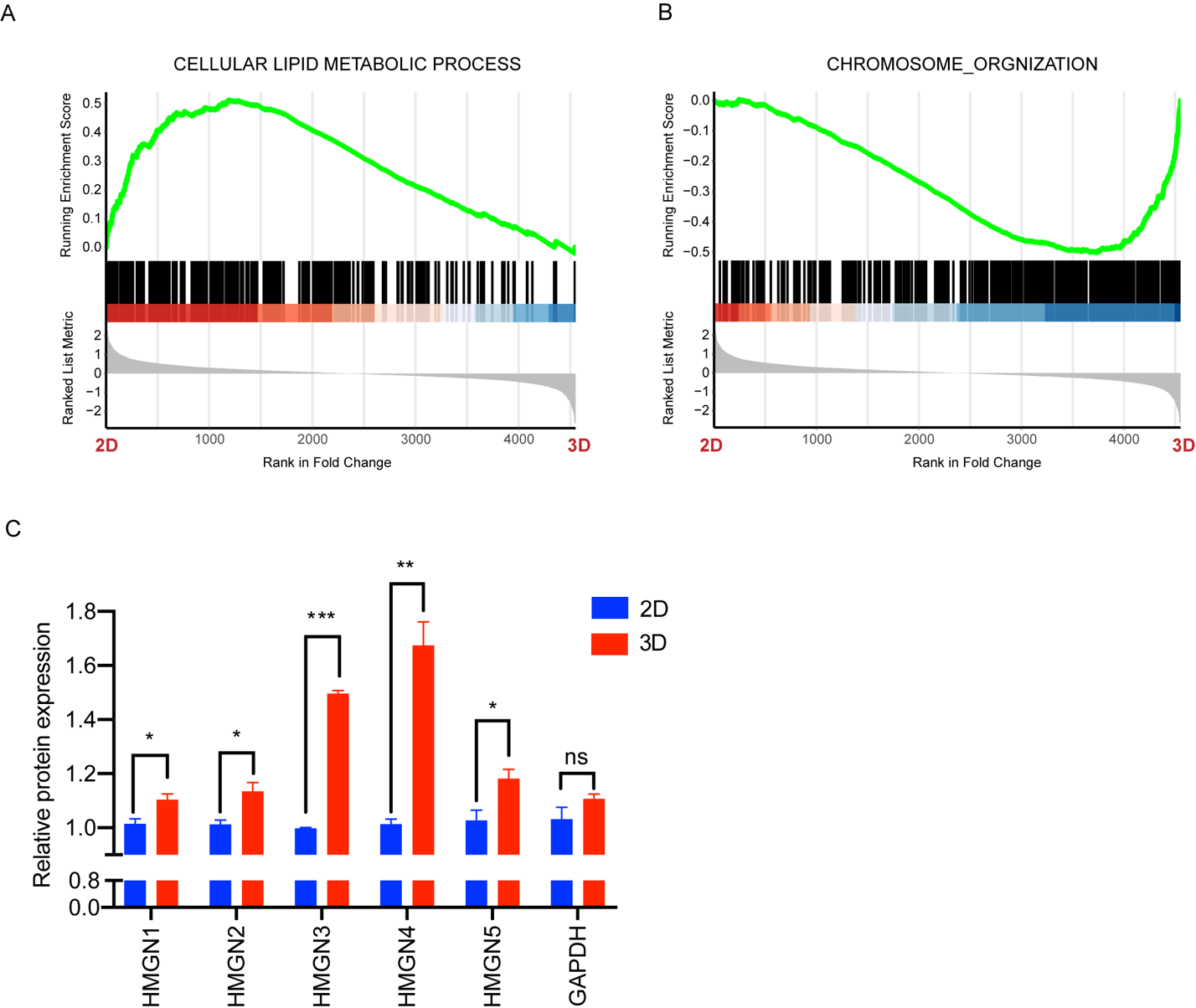

**Fig. S1. Proteomic analysis** **in adherent cells and tumorspheres of MDA-MB-231.** (A-B) GESA enrichment plot of expression change for the indicated gene sets according to the protein levels in tumorspheres (3D) vs. adherent cells (2D). (C) HMGNs abundance was detected by MS in tumorspheres and adherent cells of MDA-MB-231. Statistical significance was determined by Student's two-tailed t-test. *p <0.05, **p <0.01, ***p <0.001.

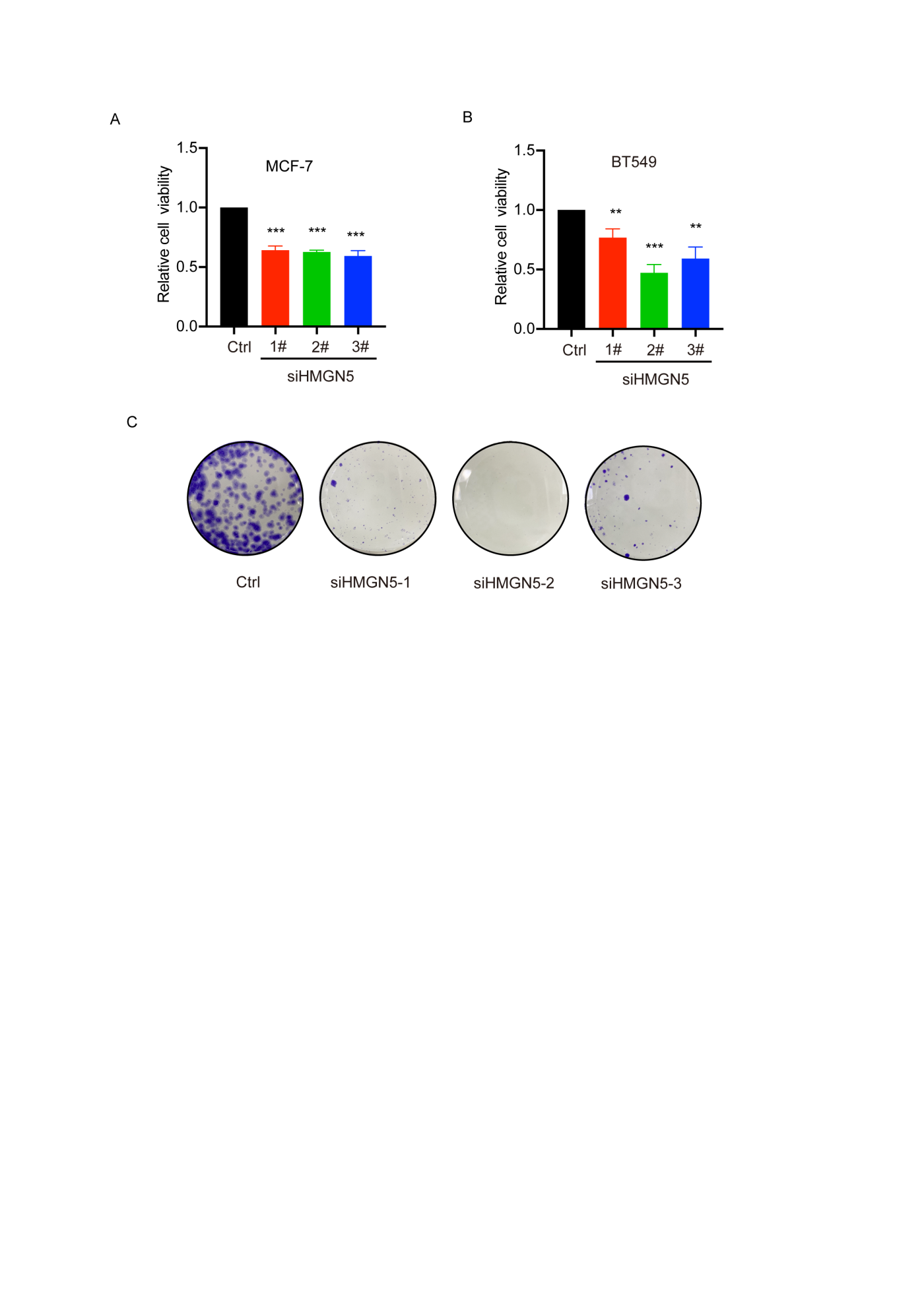

**Fig. S2. Effects of HMGN5 depletion on breast cancer cells growth and clonal expansion *in vitro*.** (A-B) Relative cell viabilities of MCF7 cells (A) and BT549 cells were determined by CCK-8 under the transient transfection of three different siHMGN5. (C) Colony formation assay in control (Ctrl) and three HMGN5 knockdown (siHMGN5) MDA-MB-231 cells. Significance was determined by one-way ANOVA test. **p <0.01, ***p <0.001 as indicated.

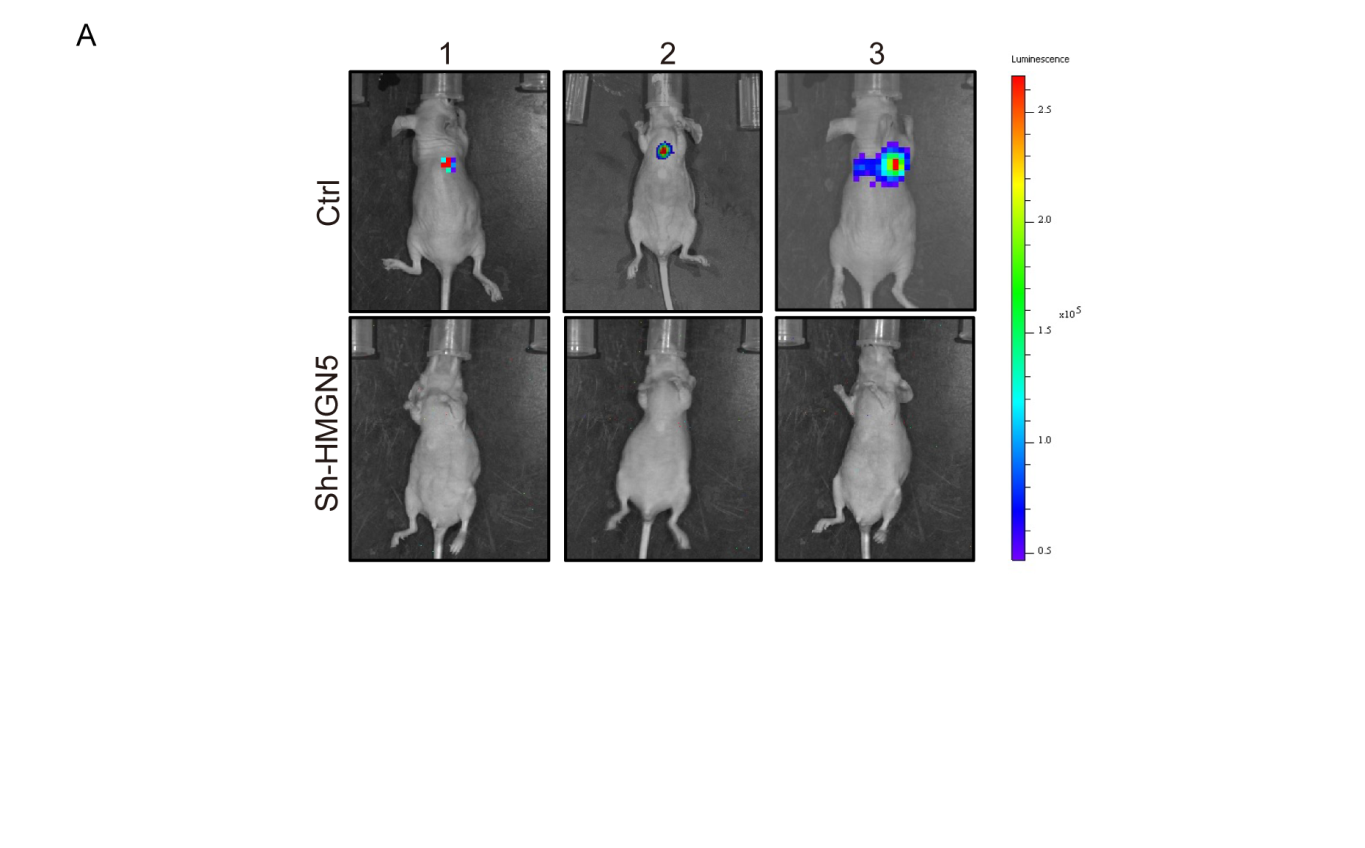

**Fig. S3. Effect of HMGN5 knockdown on metastasis *in vivo*.** (A) Bioluminescence images of three representative BALB/c nude mice in the indicated groups at 8 weeks after tail vein injections.

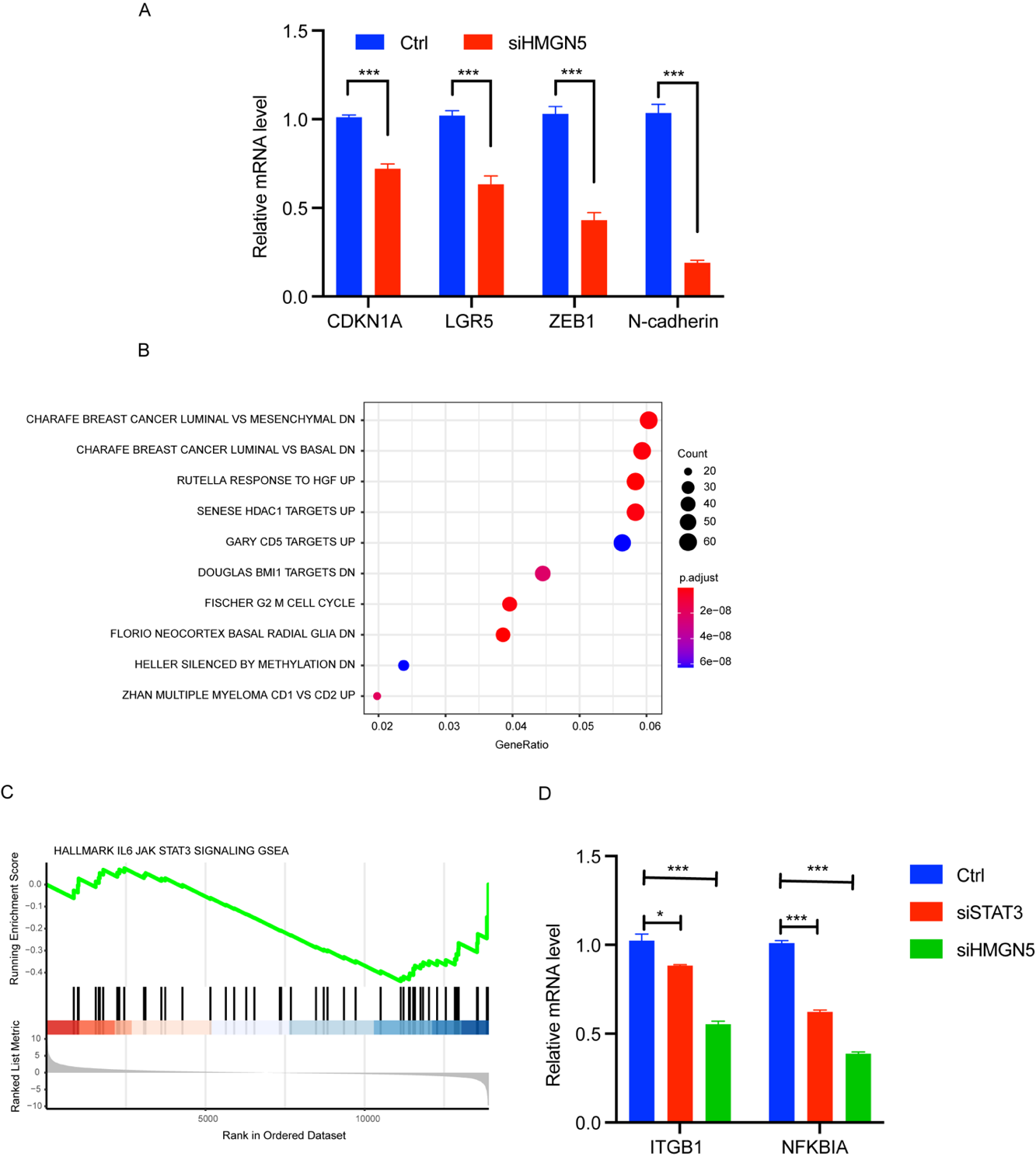

**Fig.S4. HMGN5 is involved in EMT and IL6/STAT3 signaling.** (A) CDKN1A, LGR5, ZEB1, and N-cadherin mRNA levels in MDA-MB-231 cells transfected with siHMGN5 or control siRNA are determined by RNA-Seq. (B) GSEA of curated gene sets in MsigDB based on the rank of expression foldchange in HMGN5 deficient cells vs. control cells. (C) GESA enrichment plot of expression change for the indicated gene sets according to the expression levels in HMGN5 knockdown (siHMGN5) and control MDA-MB-231 cells. (D) ITGB1 and NFKBIA mRNA levels in HMGN5 knockdown (siHMGN5), STAT3 knockdown (siSTAT3) and control MDA-MB-231 cells. Significance was determined by one-way ANOVA test. *p <0.05, **p <0.01, ***p <0.001

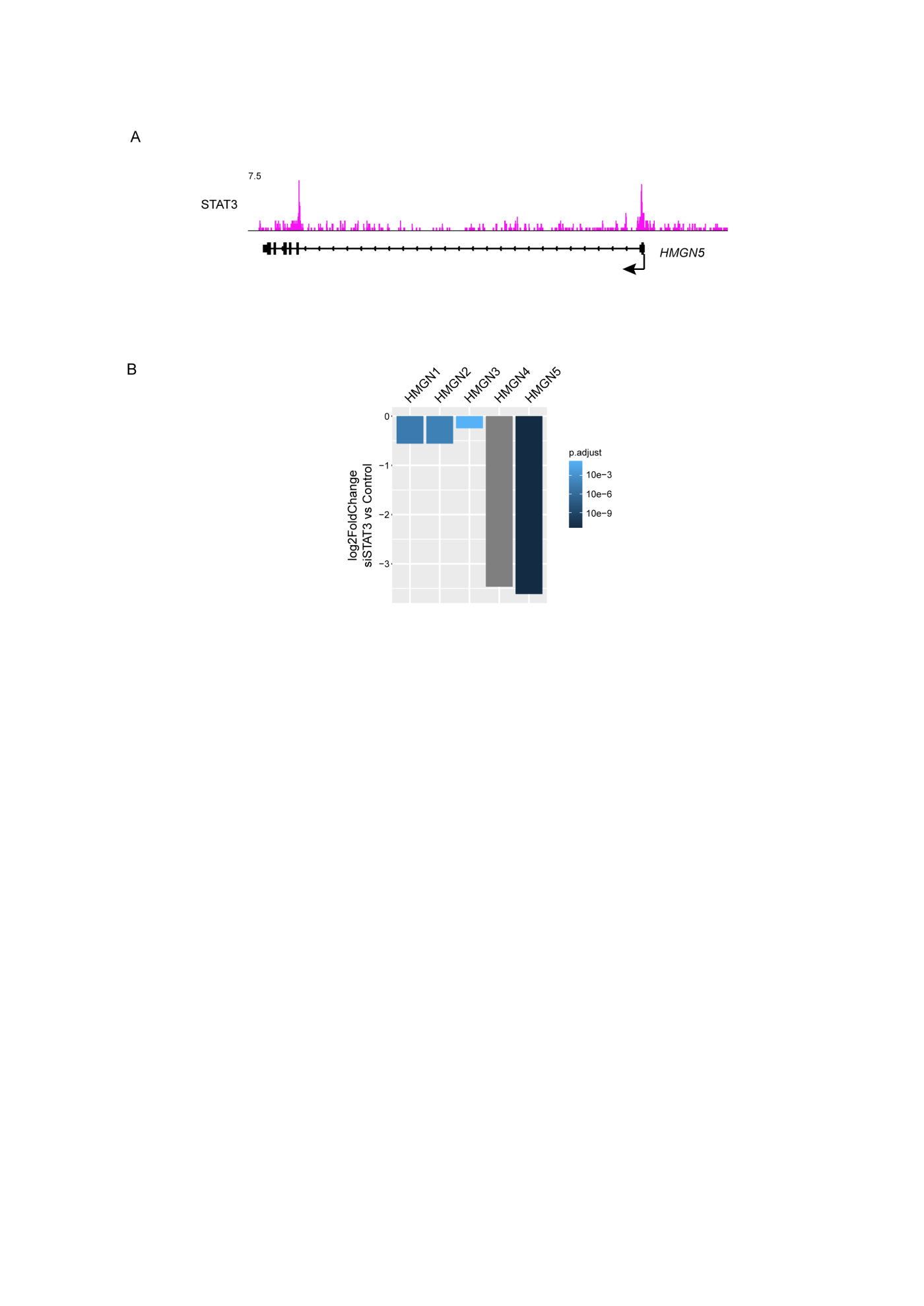

**Fig.S5. STAT3 controls the transcription of HMGN5 via binding to its promoter.** (A) STAT3 binding profile at *HMGN5* *locus*. (B) The differential expression changes of HMGNs in STAT3-deficient MDA-MB-231 vs. control cells are determined by RNA-Seq.

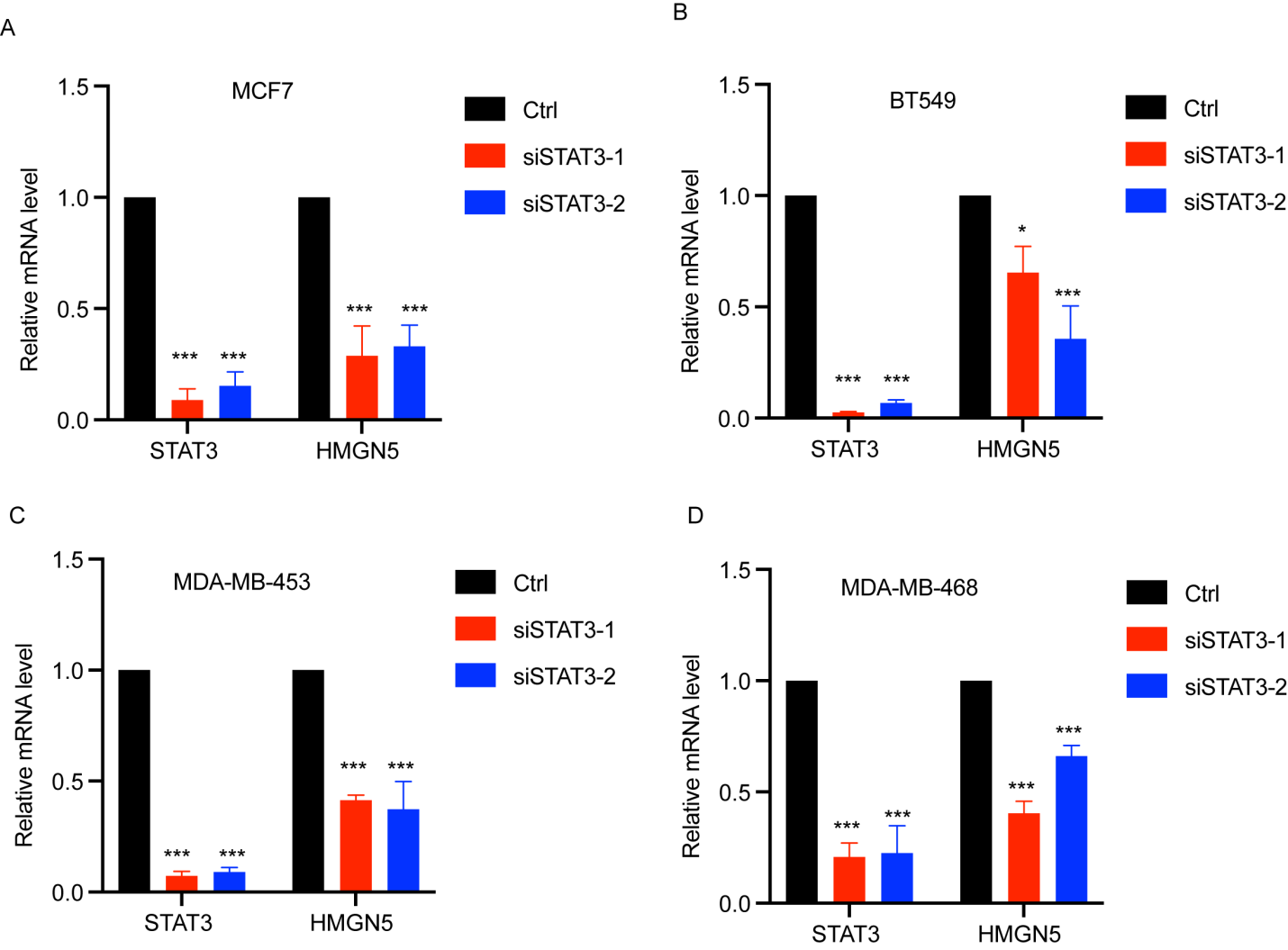

**Fig.S6. STAT3 universally regulates HMGN5 expression in breast cancer cells.** (A-D) STAT3 and HMGN5 mRNA levels in the indicated breast cancer cell lines transfected with two different STAT3 siRNAs are determined by RT-qPCR.

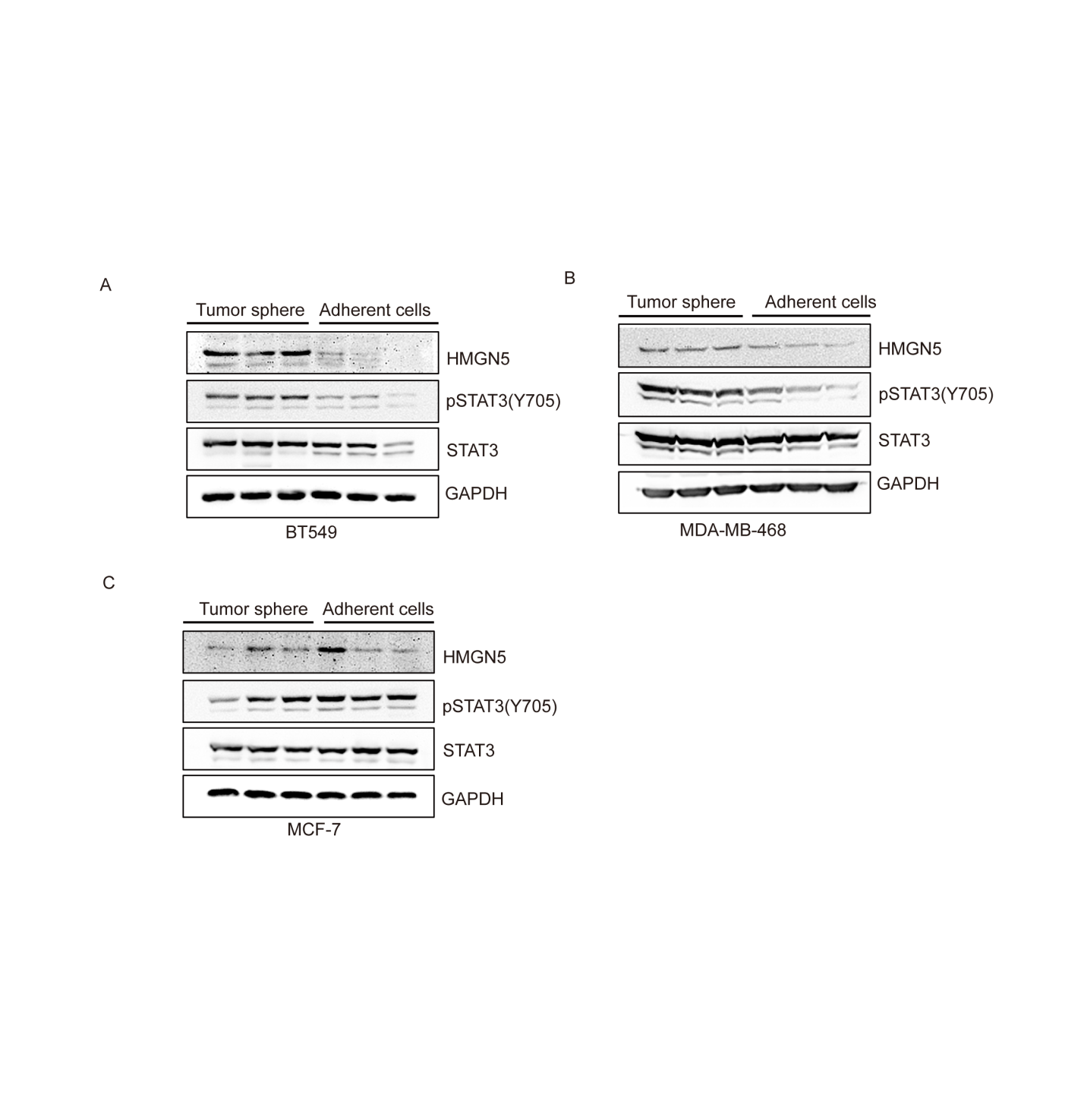

**Fig.S7. The simultaneous increase of p-STAT3 and HMGN5 in 3D-cultured breast cancer cells is in a cell type-specific manner.** (A-C) HMGN5 and p-STAT3 levels were detected by western blotting in 2D- and 3D-cultured breast cancer cells.

Table S1.

Differentially-expressed proteins in adherent cells versus tumorspheres of MDA-MB-231 (n=3) detected by Mass spectrometry (MS).

Table S2.

Primer Sequences for RT-qPCR Analysis

| Gene | Direction | Sequence (5’-3’) |
| --- | --- | --- |
| GAPDH | Forward | GGAGCGAGATCCCTCCAAAAT |
|  | Reverse | GGCTGTTGTCATACTTCTCATGG |
| STAT3 | Forward | CAGCAGCTTGACACACGGTA |
|  | Reverse | AAACACCAAAGTGGCATGTGA |
| HMGN1 | Forward | GCGAAGCCGAAAAAGGCAG |
|  | Reverse | TCCGCAGGTAAGTCTTCTTTAGT |
| HMGN2 | Forward | AGTTTGGCTTGGAATGCTGC |
|  | Reverse | AGCAGAACGTACCCTGTTCC |
| HMGN3 | Forward | GAGCCCACAAGACGGTCTG |
|  | Reverse | TCTTCCCTTTAGCACCTCTGC |
| HMGN4 | Forward | GATCAGCTCGGTTGTCTGCTA |
|  | Reverse | GCAGGGTTGTTCCCATCCTT |
| HMGN5 | Forward | CAGGTCAAGGTGATATGAGGCA |
|  | Reverse | GCTTGGGCACTTGTATCTATGT |

NOTE. Glyceraldehyde-3-phosphate dehydrogenase (GAPDH) was amplified as an internal control.

Table S3.

**Oligonucleotide sequences targeting STAT3 and HMGN5 mRNA**

| Target gene |  | Sequence (5’-3’) |
| --- | --- | --- |
| STAT3 | siRNA 1# | CCGUGGAACCAUACACAAATT |
|  | siRNA 2# | CCCGGAAAUUUAACAUUCUTT |
|  | siRNA 3# | GGGACCUGGUGUGAAUUAUTT |
|  | shRNA | CGGATCATAAGGTCAGGAGAT |
| HMGN5 | siRNA 1# | GCAAGAAGCAGUUGUUGAATT |
|  | siRNA 2# | GCUUGUGCCAGUUACACCATT |
|  | siRNA 3# | GCCCAAGCAGUUGCUGAAATT |
|  | shRNA | CACAGCCTTTCTTTAGCATAT |
